# MONTAGE: A Computation Framework to Identify Spatially Resolved Functional Enrichment Gradients in the Tissue Microenvironment via Spatial Communities

**DOI:** 10.1101/2025.04.30.651486

**Authors:** Weiruo Zhang, Zinaida Good, Marc A. Baertsch, Guolan Lu, John W. Hickey, Rachel Hildebrand, Serena Chang, Andrew J. Gentles, John B. Sunwoo, Quynh-Thu Le, Christina S. Kong, Garry P. Nolan, Sylvia K. Plevritis

**Author notes:** **Correspondence to**: Sylvia K. Plevritis, PhD, William M. Hume Professor in the School of Medicine Professor of Biomedical Data Science and Radiology Chair, Department of Biomedical Data Science Stanford University, School of Medicine.

## Abstract

Single-cell spatial-omics has advanced our understanding of how tumor microenvironment contributes to cancer progression. However, most single-cell spatial-omics studies focus on cell types and neighborhoods, offering limited information about functional and clinical relevance of cellular organization. We introduce MONTAGE, a computational framework to reconstruct, functionally analyze, and identify clinically relevant cellular spatial communities (SCs). MONTAGE generates a gene signature matrix of SCs from integrated atlases that offer single-cell resolution with transcriptomics, and reconstructs tissue maps (“montages”), reflecting sequential SC compositional changes along spatial gradients of biological function enrichments. The MONTAGE signature matrix can also be used to deconvolve bulk and spot-based spatial transcriptomic data, and combining with clinical metadata, allows assessing SC clinical relevance. In a head and neck cancer study, MONTAGE reveals SCs enriched with malignant cells and granulocytes shifting to SCs enriched with macrophages and fibroblasts along gradient of epithelial-mesenchymal transition, and clinical prognostic significance of the SCs.

## Introduction

Solid tumors are ecosystems consisting of cancer cells that accumulate mutations in coordination with non-cancer cells (the tumor microenvironment, TME), where the proximity, and thereby likely interactions, of cancer cells, immune cells, and stromal cells contribute to malignancy and drug resistance ^1–3^. Advances in single-cell spatial-omics that provide molecular profiles and spatial locations of individual cells have profoundly enhanced our analysis of TME cellular spatial organization ^4,5^. Yet, even more valuable is the ability to recognize and monitor heterogenous TME cell environments as dynamic spatial communities (SCs), to better understand tissue organization, disease mechanisms, and potential therapeutic targets. Single-cell spatial-omics studies are limited by molecular multiplexing that constrain the ability to comprehensively characterize cell types and states in their functional context ^6,7^. Moreover, due to cost considerations, single-cell spatial-omics studies often rely on use of tissue microarrays (TMA) constructed from small tissue cores (typically < 1mm in diameter). These samples typically focus only on cancer-enriched regions, limiting the ability to interrogate the role of immune and stromal cells in cancer properties and progression. In contrast, use of whole-slide tissue sections provide a larger view of the TME, enabling analysis of its many cancer and non-cancer cell types. Currently, whole-slide spatial-omics studies at single-cell resolution typically include fewer than 20 samples with moderately multiplexed spatial proteomic features (< 100 markers). Although several methods are available to integrate these spatial proteomic and single-cell RNA sequencing (scRNA-seq) data from adjacent tissue ^8–10^, few approaches can use this information to identify spatially resolved gene enrichments that underly functional biological changes, including those with clinical relevance.

To overcome these limitations, we present MONTAGE, a computational framework that leverages typical integrated tissue atlases generated from whole-slide spatial proteomics and scRNA-seq data, and can be generalized to single-cell transcriptomic datasets as they accrue. Using the integrated atlases, MONTAGE reconstructs, functionally analyzes, and identifies clinically relevant TME spatial communities (SCs). To achieve this, MONTAGE first generates a gene signature matrix of SCs and then derives a sequence of SC compositions (“montages”) along gradients of functional gene enrichment. Such gradients reflect increased presence or activity of specific biological pathways, cellular functions, or molecular interactions that contribute to cancer progression, immune response, or drug resistance. MONTAGE thus illuminates the dynamic evolution of biological processes, including the progressive acquisition of increasingly malignant characteristics within the TME.

MONTAGE also provides an alternative to common deconvolution methods for bulk transcriptomics ^11–13^ and region/spot-based spatial transcriptomics data ^14–16^ . Different from existing deconvolution methods that deconvolve cell type compositions, MONTAGE allows deconvolving transcriptomic datasets into spatial community compositions. MONTAGE is also different from methods like EcoTyper ^17^ that estimate cellular community compositions from co-expression patterns across bulk gene expression profiles, because MONTAGE cellular spatial communities were directly identified from single-cell-resolved spatial data. When used to deconvolve large-scale, publicly available transcriptomic datasets with available clinical metadata, MONTAGE enables identification of clinically relevant spatial communities.

We demonstrate the ability of MONTAGE to identify sequential SC compositional changes along functional gradients associated with increasing malignant phenotypes related to biology processes including glycolysis, interferon gamma signaling, and the epithelial-to-mesenchymal transition (EMT) within the TME of a study cohort of head and neck squamous cell carcinoma (HNSCC). These findings are associated with clinical outcomes, demonstrating that MONTAGE generates functionally and clinically valuable information about how SC compositions contribute to pathology in the TME.

## Results

### Overview of the MONTAGE computational framework and study design

The MONTAGE framework first creates a spatial community (SC) gene signature matrix by leveraging a single-cell atlas derived by integrating scRNA-seq and single-cell multiplexed imaging-based spatial proteomics data (**Fig 1A**). MONTAGE requires two input matrices. First, the matrix ***G*** is a *p* × *k* matrix that contains the average gene expressions for *p* cell type gene markers of *k* cell types and is derived from scRNA-seq data. Second, the matrix ***S*** is a *k* × *n* matrix that has the proportions of *k* cell types in each of the *n* spatial communities, derived from single-cell spatial proteomics data. The MONTAGE SC gene signature matrix ***M*** is created by multiplication of matrices ***G*** and ***S*** and encodes each SC by weighting cell type marker gene expressions with corresponding cell type proportions in each SC. Descriptions of how to construct matrices ***G****, **S**,* and ***M*** using our study cohort, and more directly from single-cell spatial transcriptomic data, are described in Methods.

**Figure 1.**
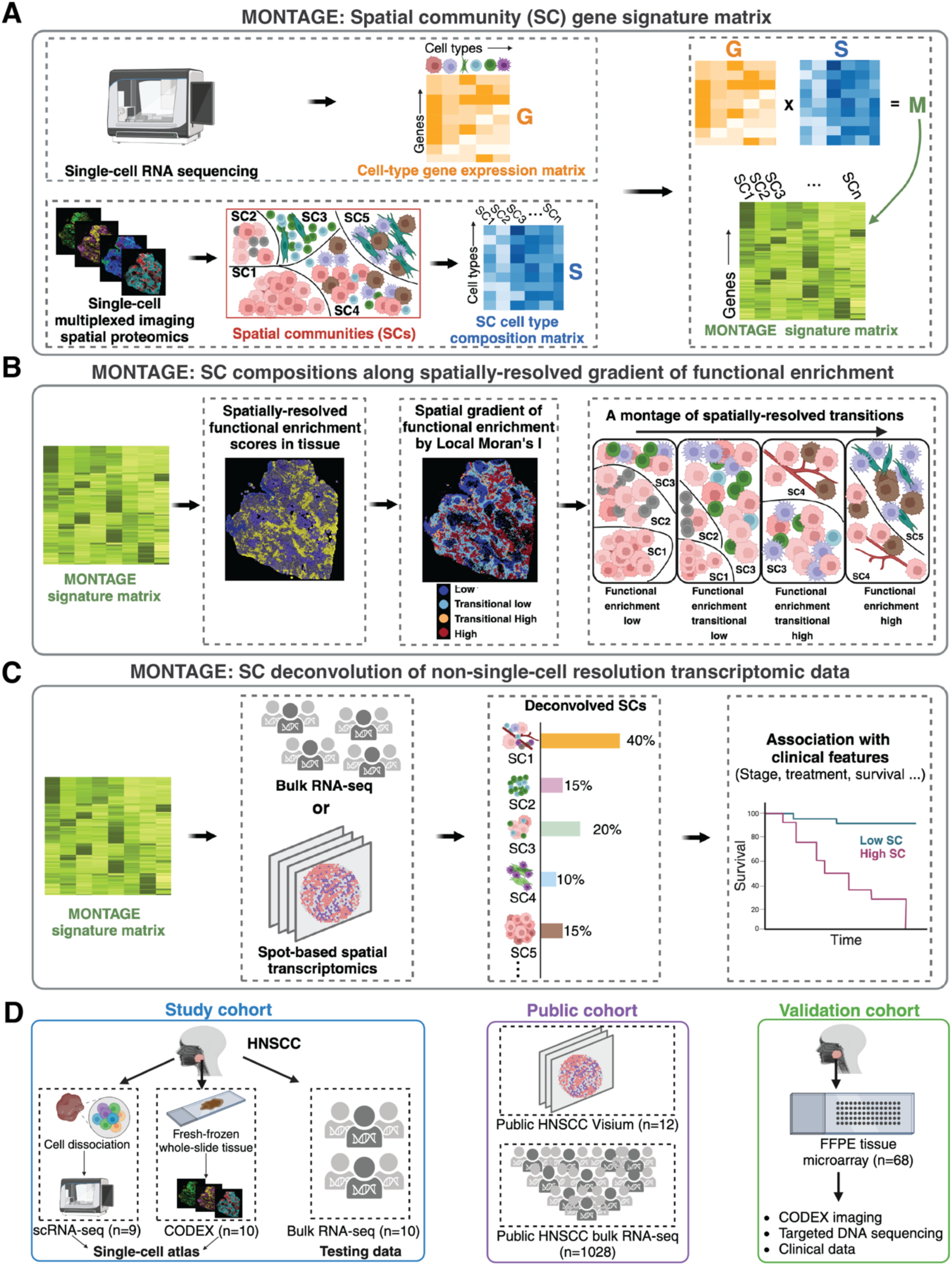
Schematic workflow of the MONTAGE framework and data overview. **(A)** The MONTAGE framework creates a spatial community (SC) gene signature matrix by integrating single-cell RNA sequencing (scRNA-seq) data and single-cell spatial proteomics data. **(B)** The MONTAGE signature matrix enables displaying gene functional enrichment in a spatial context on spatial proteomics images, identifying a spatially resolved gradient along increasing functional enrichment and creating a montage of spatial community composition changes along the spatially resolved gradient of functional enrichment. The functional enrichments can be performed using gene sets of any biological pathways, cellular functions, or drug responses. **(C)** The MONTAGE gene signature matrix enables deconvolving large-scale bulk RNA-seq or spot-based spatial transcriptomics data into SC proportions and associating the SC proportions with clinical features to assess the clinical relevance of the SCs. **(D)** An overview of the datasets used to demonstrate MONTAGE in the study of head and neck squamous cell carcinoma (HNSCC).

The MONTAGE SC gene signature matrix ***M*** can be used for functional analysis of SCs in the TME as well as to deconvolve large non-single-cell transcriptomic datasets. First, the MONTAGE signature matrix ***M*** is used to reconstruct a tissue map image with each cell assigned to its SC and a functional enrichment score (**Fig 1B**). MONTAGE applies the local Moran’s I spatial autocorrelation to identify regions of distinct functional enrichment, ranging from regions of “low” to “translational” to “high” functional enrichment. MONTAGE then creates “montages” as sequences of SC compositions along spatially resolved gradients of functional enrichment. From these montages, MONTAGE pinpoints spatially defined transitions of SC composition, in which functional pathways shift. Second, the MONTAGE signature matrix ***M*** enables the deconvolution of large external cohorts of transcriptomics data, such as those from The Cancer Genome Atlas (TCGA), leveraging existing deconvolution methods, such as CIBERSORT ^11^, EPIC ^12^, or xCell ^13^ but replacing traditional cell type-only deconvolution with SC deconvolution and then associating SC compositions with clinical features (**Fig 1C**).

We applied MONTAGE to a study cohort of 8 HNSCC patients, from whom we acquired primary and metastatic lymph node tissue samples. These included 9 tissue samples profiled with scRNA-seq, 10 samples profiled with whole-slide CODEX ^18^ imaging spatial transcriptomics, and 10 samples profiled with bulk RNA sequencing (RNA-seq) from proximal tissue sections (**Fig 1D**). A summary of patient clinical information was provided in Supplementary Table S1.

We applied MONTAGE to deconvolve a public cohort of a Visium spatial transcriptomics dataset of 12 HNSCC clinical samples. We also curated, publicly available bulk RNA-seq datasets from HNSCC clinical samples (N =1,028) with survival outcome data for assessing clinical significance of MONTAGE deconvolved spatial communities (**Fig 1D**). Finally, we established an independent cohort of 68 HNSCC patients by constructing a tissue microarray (TMA) of tumor-enriched regions, toward validating MONTAGE-derived findings from our study cohort, specifically within tumor-enriched areas (**Fig 1D**). For our HNSCC study cohort, we first created an integrated atlas of 483,646 single cells from CODEX spatial proteomic data and 21,496 single cells from scRNA-seq data (**Fig 2A-B**). The cell types were annotated based on Seurat ^8^ clustering using scRNA-seq (**Supplementary Fig S1A-H**) and matrix ***G*** was constructed with the gene expressions from cell type marker. We then assigned cell types in the CODEX data (**Supplementary Fig S1I, S2, S3A**) using the transferring learning function in Seurat (see **Methods** section).

**Figure 2.**
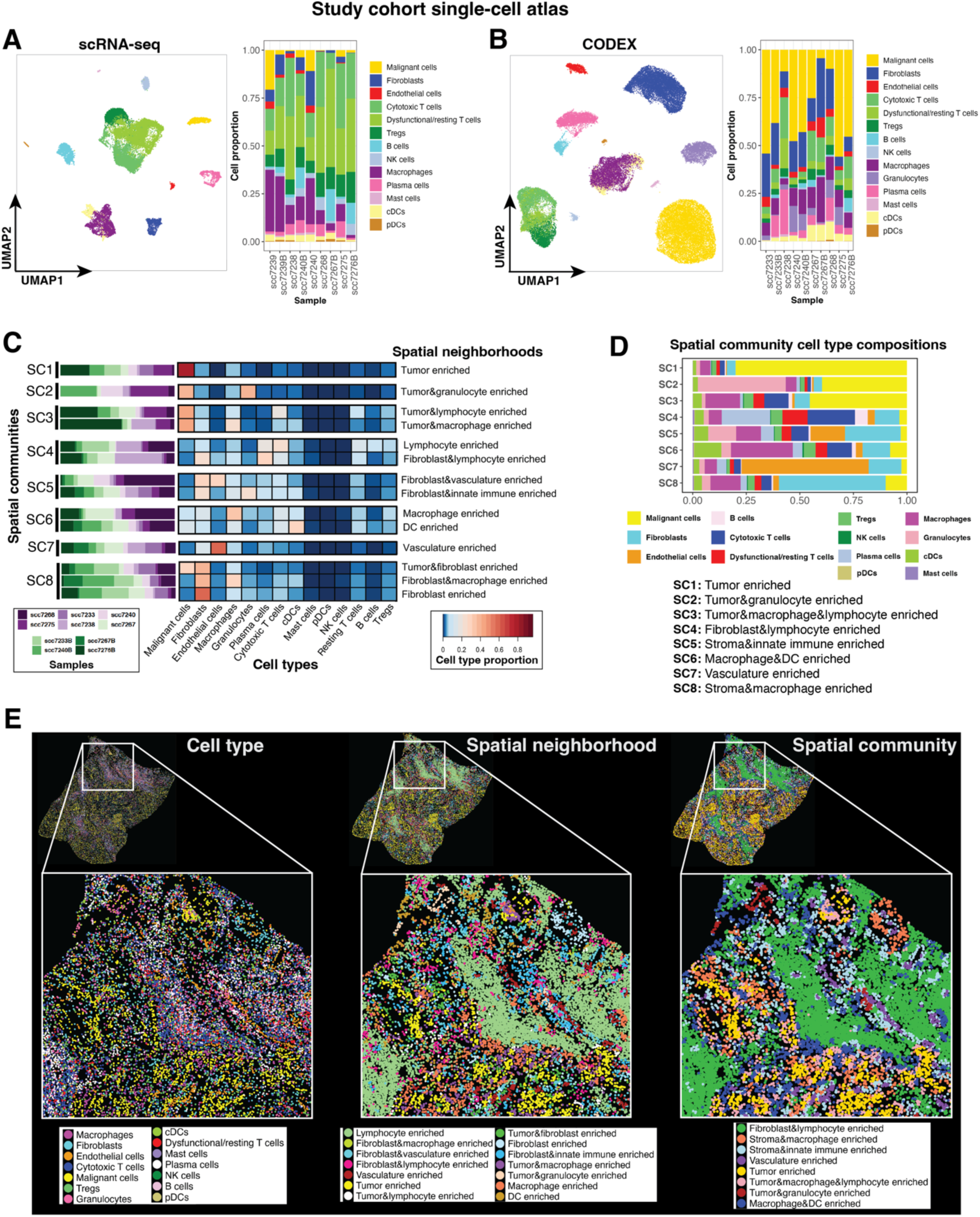
Overview of the cell types identified in the study cohort single-cell atlas and MONTAGE spatial community (SC) identification. **(A)** Uniform manifold approximation and projections (UMAPs) of scRNA-seq and barplot showing cell type proportions across samples in the scRNA-seq data of the HNSCC single-cell atlas from the study cohort. **(B)** UMAPs of spatial proteomics (CODEX) cells and barplot showing cell type proportions across samples in the spatial proteomics CODEX data of the HNSCC single-cell atlas from the study cohort. 10% of sampled cells were used in plotting the UMAP for visualization. **(C)** Heatmap shows the cell type proportions in the 14 spatial neighborhoods that clustered into 8 spatial communities (SCs), identified using the CODEX data from the HNSCC study cohort single-cell atlas. The barchart on the left of the heatmap displays the distribution of patient samples within each spatial neighborhood. **(D)** Barplot showing the cell type compositions in each SC, denoted as SC1 to SC8 and named by major cell type constitutions. **(E)** A representative whole-slide tissue sample in the HNSCC study cohort plotted with identified cell types (left panel), spatial neighborhoods (middle panel), and spatial communities (right panel). Each plotted dot represents a cell centroid.

### MONTAGE identifies HNSCC SCs and their gene signatures

MONTAGE identifies spatial communities (SCs) within tissue using an adaptation of common strategies ^19,20^. We first performed cell-based nearest-neighborhood analysis on the CODEX images of our HNSCC study cohort (see **Methods** section) to quantify densities of cells in local proximity and identified 14 spatial neighborhoods displaying distinct local cell-density patterns (**Fig 2C**). Clustering the cell-type compositions in the 14 spatial neighborhoods, we obtained 8 spatial communities (denoted SC1, SC2, … SC8) (**Fig 2C**). Compared to the spatial neighborhoods, SCs are larger tissue regions appearing more consistently across samples. Each SC is annotated with its dominant cell types (**Fig 2D, Supplementary Fig S3B**). We determined the number of SCs based on the best similarity score between paired testing bulk RNA-seq data and CODEX data in our HNSCC study cohort (**Supplementary Fig S3C,** Methods). To evaluate the robustness of the SCs, we performed a leave-out-sample analysis (**Supplementary Fig S3D**), which demonstrated that SC1, SC2, SC3, SC4, SC7, and SC8 could be robustly identified within more than five samples; SC5 and SC6 exhibited lower cross-sample robustness, potentially due to higher cell-type heterogeneity compared to other SCs. A representative sample of a primary HNSCC tumor from a patient with lymph node metastasis (N+) is shown in **Fig 2E** to illustrate cell types and enrichment within spatial neighborhoods and SCs. A sample from a primary tumor of a patient without lymph node metastasis (N0) and another tumor from a metastatic lymph node are shown in **Supplementary Fig S4A-B**. SCs could be further subdivided into three classes (**Supplementary Fig S4C**). To associate various levels of tissue structure, we built a hierarchical network graph ^20^ with each level of the graph displaying structures with reduced granularity (**Supplementary Fig S4D**).

Matrix ***G,*** created from scRNA-seq data, together with matrix ***S,*** created from the CODEX imaging data, represent 14 cell types in 8 SCs and are used to build the MONTAGE signature matrix ***M*** (**Fig 3A, Supplementary Table S2**). To verify the signature matrix ***M***, we applied it to deconvolve paired bulk RNA-seq data in our study cohort. We observed a high correlation between deconvolved SC compositions using the MONTAGE signature matrix and those obtained from CODEX images directly (**Fig 3B**). Within-sample comparisons also showed that deconvolved bulk RNA-seq SC compositions generated high similarity scores to CODEX SC compositions across each of the 10 HNSCC samples (**Fig 3C**). We performed gene-set enrichment analysis ^21^ on each SC gene signatures to functionally characterize SCs (**Fig 3D**) using Hallmark genesets from MsigDB ^22^. Of note, SCs with dominant proportions of tumor cells (SC1, SC2) were functionally enriched for the P53 pathway, glycolysis, and cell proliferation.

**Figure 3.**
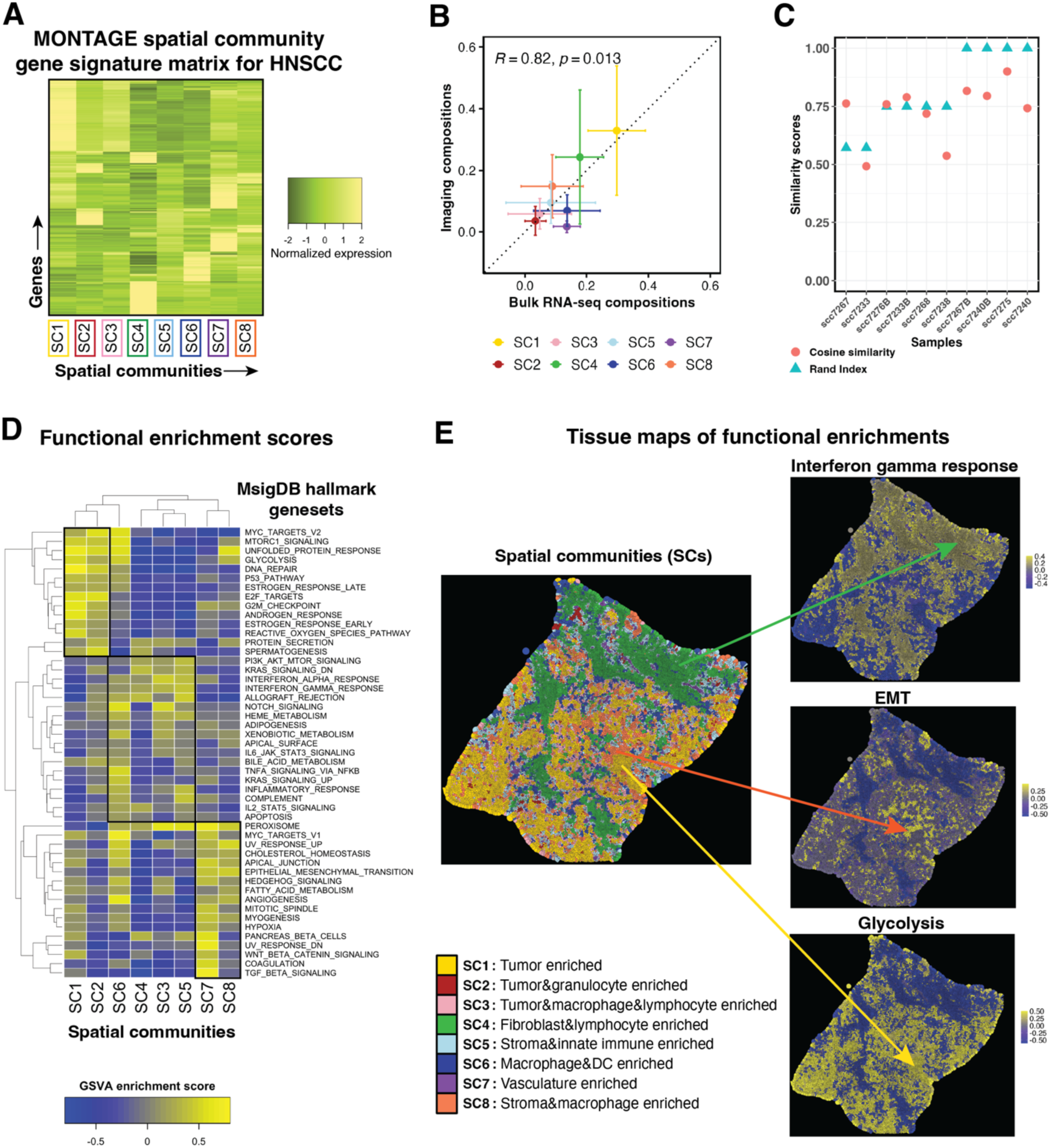
MONTAGE spatial community (SC) gene signature matrix for HNSCC study cohort and tissue maps of functional enrichments. **(A)** Heatmap displaying the normalized expressions of the MONTAGE signature matrix created from the HNSCC study cohort. **(B)** Comparison of SC compositions derived from bulk RNA-seq deconvolution using the MONTAGE signature matrix and the SC compositions obtained from the CODEX imaging data of the HNSCC study cohort shows high correlation. Bulk RNA-seq and CODEX imaging data were generated from proximal slices. **(C)** Similarity scores for the deconvolved SC proportions of bulk RNA-seq and imaging-derived SC proportions demonstrate high concordance across all the clinical samples in the HNSCC study cohort. **(D)** MONTAGE signature matrix enabled functional enrichment scores using the MsigDB Hallmark genesets for each spatial community (SC). **(E)** Tissue maps of spatially resolved functional enrichment scores are created by assigning each cell in a whole-slide spatial proteomics image with its respective functional enrichment score, based on the specific spatial community to which the cell belongs. Enrichments of interferon response, epithelial-mesenchymal transition (EMT) and glycolysis are used for illustration purposes.

SCs with large proportions of immune cells (SC3, SC4, SC5, SC6) were enriched for an interferon response. SCs with large proportions of stromal cells (SC7, SC8) were enriched for biological functions including angiogenesis, the epithelial-mesenchymal transition (EMT), and hypoxia ^23^. After assigning each cell with its corresponding SC functional enrichment score, MONTAGE superimposed these scores onto cell-segmented tissue images, thereby producing a tissue map for functional enrichment of such processes such as interferon gamma, EMT and glycolysis (**Fig 3E, Supplementary Fig S5A**). In this manner, MONTAGE enables visualization of functional enrichment within the TME in a spatial context at single-cell resolution.

### MONTAGE derives a montage of SCs along a spatially resolved gradient of functional enrichment

MONTAGE applies the local Moran’s I ^24^ to identify spatially resolved regions with distinct functional enrichment scores and detects spatial gradients linked to functional enrichment shifts. Because EMT has been associated with metastasis for multiple tumor types, including HNSCC^25,26^, we first focused on the identification and analysis of spatial gradient associated with EMT functional enrichment. MONTAGE applied the local Moran’s I on spatially resolved functional enrichment EMT scores (**Fig 4A**) and categorized four spatially resolved regions along a gradient of increasing EMT-enrichment scores: i) low-EMT, ii) transitional-low EMT, iii) transitional-high EMT, and iv) high-EMT regions. These data were used to calculate average SC compositions for each of the four EMT regions across the 10 CODEX images (**Fig 4B**), revealing characteristic patterns of SCs and EMT-enrichment (**Fig 4C**). With this montage of spatially resolved EMT **(Fig 4D**), we identified that in the low-EMT region of the HNSCC TME, the most abundant SCs were tumor-enriched (SC1), tumor- and granulocyte-enriched (SC2), and lymphocyte-enriched (SC4), suggesting that epithelial tumor cells are mostly co-located with granulocytes and lymphocytes in low-EMT regions. In transitional-EMT regions of the HNSCC TME, compositions of SC1, SC2, and SC4 (abundant with lymphocytes) decreased, whereas SCs with large proportions of innate immune cells such as macrophages and dendritic cells (SC3, SC5, SC6) increased. In high-EMT regions of the HNSCC TME, SCs enriched with macrophages and cells of the vasculature and stroma (SC7, SC8) were predominant, suggesting a role for stromal cells and macrophages in promoting the EMT ^27^ (**Fig 4D**).

**Figure 4.**
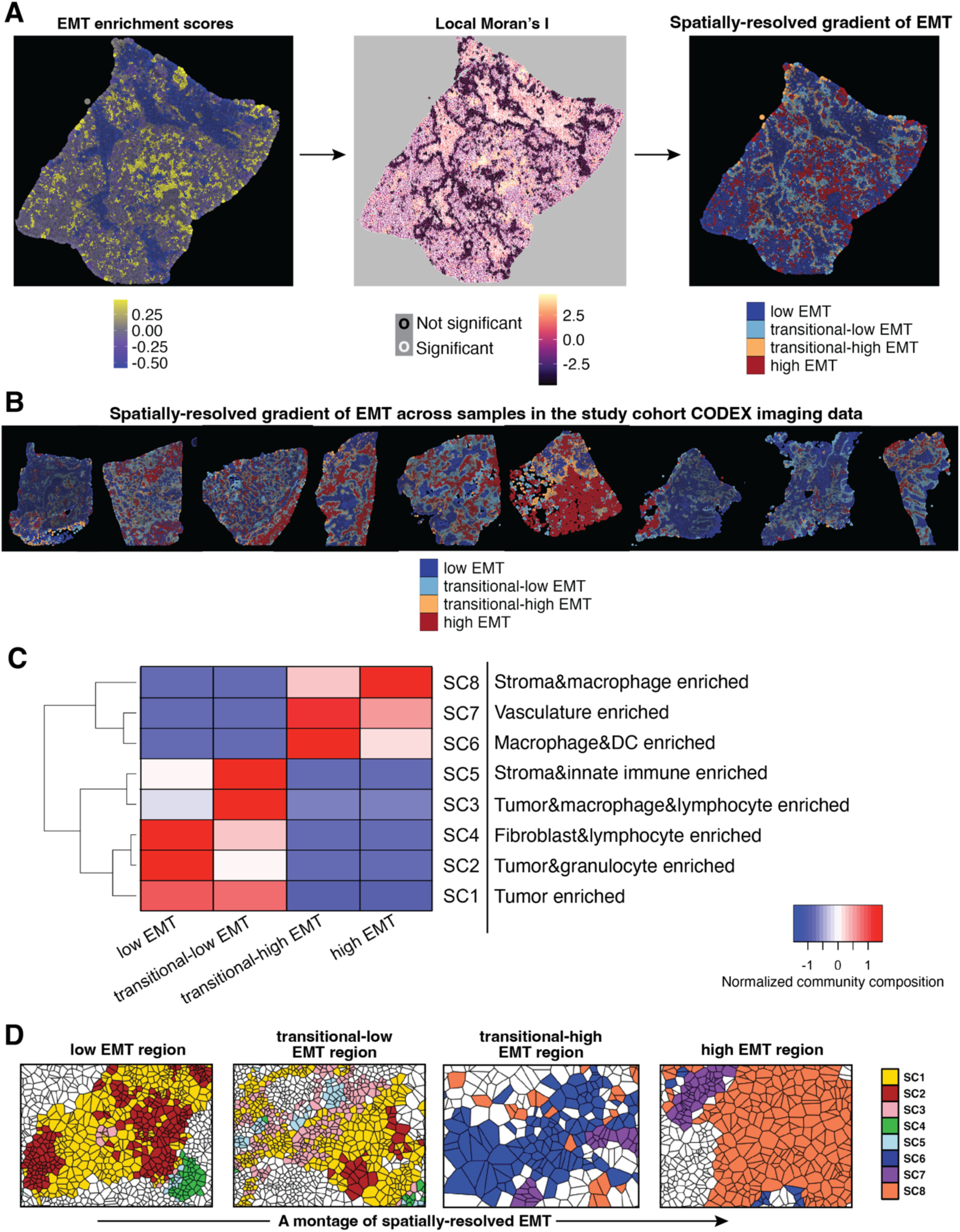
MONTAGE creates a montage of spatial community (SC) compositions along spatially resolved gradient of gene functional enrichment of EMT. **(A)** Applying local Moran’s I statistics to derive spatially resolved gradient along increasing epithelial-mesenchymal transition (EMT) functional enrichment scores, illustrated on a representative sample in the HNSCC study cohort. **(B)** Identified regions along spatially resolved gradient of increasing EMT enrichment of all other whole-slide tissue samples in the study cohort. **(C)** Heatmap showing normalized SC composition scores progression along the spatially resolved gradient of EMT aggregated across 10 spatial proteomics whole-slide tissue samples in the study cohort. **(D)** Voronoi graphs of representative regions color coded by SCs along spatially resolved gradient of EMT; collection of images represents a montage for cellular reorganization along a spatially resolved functional gradient of EMT.

To further illustrate this use of MONTAGE to resolve SC compositions along functional gradients in the TME, we applied MONTAGE to analyze glycolytic activity (**Supplementary Fig S5B,C**), another hallmark of tumor tissue ^28^. We observed that a lymphocytes-enriched SC was most prevalent in a low glycolytic activity region; and SCs enriched with innate immune cells and tumor cells infiltrated with lymphocytes were abundant in a transitional-low glycolytic activity region, consistent with current literature suggesting the role of lymphocytes in suppressing glycolysis ^29^. Alternatively, SCs enriched with stromal, macrophage, and dendritic cells were abundant in transitional-high glycolytic activity regions, and tumor cell-enriched SCs were most prevalent in high-glycolytic activity regions. Interestingly, compared with MONTAGE-derived EMT results, SCs with the highest metastatic potential (high-EMT) regions did not co-locate with those with highest glycolytic activity, consistent with current literature that highlights the complex relationship between glycolysis and EMT ^30^. These findings demonstrate that MONTAGE identifies cellular reorganization associated with a multiplicity of functional properties in localized regions of the TME, providing additional avenues for mechanistic investigation.

### Validation of MONTAGE HNSCC findings of SC compositional changes along spatially resolved functional gradients on an independent cohort

We validated MONTAGE-derived findings from our HNSCC study cohort by applying the MONTAGE signature matrix to deconvolve an independent, publicly available Visium dataset ^31^ consisting of 12 whole-slide HNSCC tissue samples that include pathologist annotations (**Fig 5A, Supplementary Fig S6A**). We first compared the pathologist’s annotations with the deconvolved MONTAGE SCs by grouping the deconvolved SC compositions into three spatial community classes (**Supplementary Fig S4C**). Our data reveal that the deconvolved SCs were consistent with the pathologist’s assessments (**Fig 5B, Supplementary Fig S6B-C**).

**Figure 5.**
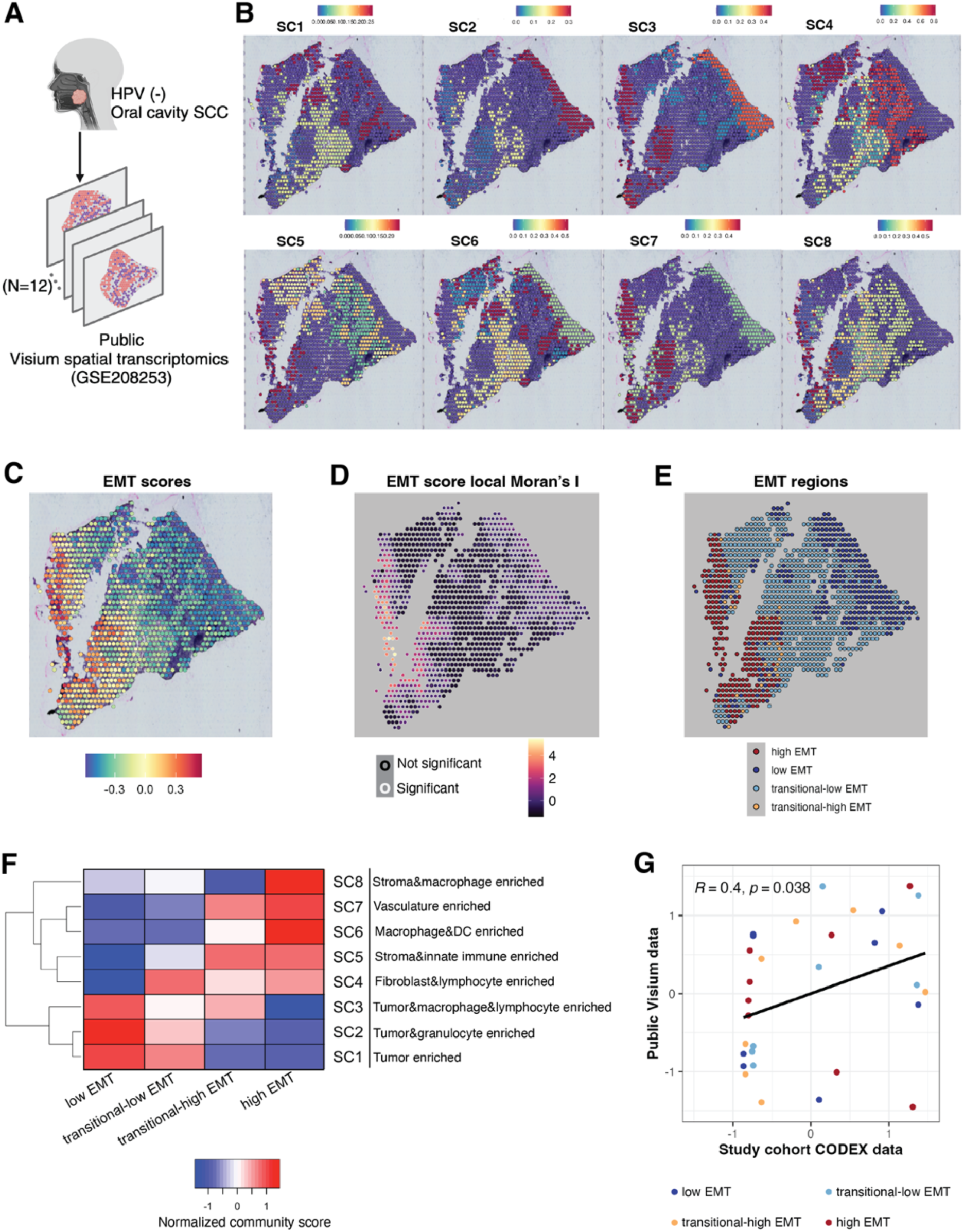
MONTAGE signature matrix used to deconvolve a public Visium spatial transcriptomics HNSCC dataset validates findings from the HNSCC study cohort. **(A)** Overview of the public spatial transcriptomic Visium HNSCC dataset (n=12). **(B)** Deconvolved MONTAGE spatial community (SC) compositions on a representative Visium whole-slide tissue sample. **(C)** Functional enrichment scores of EMT of a representative sample. **(D)** Local Moran’s I statistics of EMT enrichment scores on a representative sample. **(E)** Identified regions along spatially resolved gradient of increasing EMT enrichment on a representative sample. **(F)** Heatmap showing SC composition changes along the spatially resolved gradient of EMT aggregated across the 12 samples in the public Visium dataset. **(G)** Comparison of the normalized community scores in the EMT regions from the public Visium HNSCC data and from the CODEX data in our HNSCC study cohort showed significant correlation despite of different resolutions (Visium is spot-based, and CODEX is single-cell based).

Deconvolved SC8 (the stromal- and macrophage-enriched SC) had significantly higher compositions in HNSCC patients with lymph node metastasis (N+) compared with node-negative patients (N0) (**Supplementary Fig S6D**), further highlighting a potential role for stromal cells and macrophages to promote metastasis.

We next applied MONTAGE to identify spatially resolved EMT gradients within the Visium HNSCC dataset (see **Methods** section). MONTAGE calculated an EMT enrichment score (**Fig 5C**) and local Moran’s I (**Fig 5D**) for each spot and identified four regions along the EMT gradient characteristic of cancer progression (**Fig 5E, Supplementary Fig S6E**). By calculating SC compositions across different EMT regions and aggregating results for the 12 samples (**Fig 5F**), we determined that the Visium HNSCC data showed highly similar and statistically significant patterns of SC ensembles along the spatially resolved EMT gradient derived from our study cohort analyzed with an integrated atlas of CODEX and scRNA-seq data (**Fig 5G**). This finding demonstrates the robustness of MONTAGE, showing consistency across two independent datasets with different resolutions (Visium is spot-based, and CODEX is single-cell based).

### MONTAGE-derived HNSCC spatial community compositions are prognostically significant in deconvolved bulk transcriptomics

We applied the MONTAGE signature matrix to deconvolve bulk transcriptomic HNSCC profiles into their constituent SC compositions, leveraging publicly available data of 1,028 tumor samples from which bulk RNA-seq and survival outcomes were available (**Fig 6A**, **Supplementary Table S3**). We obtained similarity scores of ≥ 0.7 when comparing SCs derived from study cohort CODEX data to other datasets (**Supplementary Fig S7A**). In particular, HNSCC bulk RNA-seq data from The Cancer Genome Atlas (TCGA: https://www.cancer.gov/tcga) produced a similarity score of > 0.9. These findings demonstrate that MONTAGE enables deconvolving large-scale cohorts of bulk gene expression into SC compositions consistent with SCs derived directly from our study cohort.

**Figure 6.**
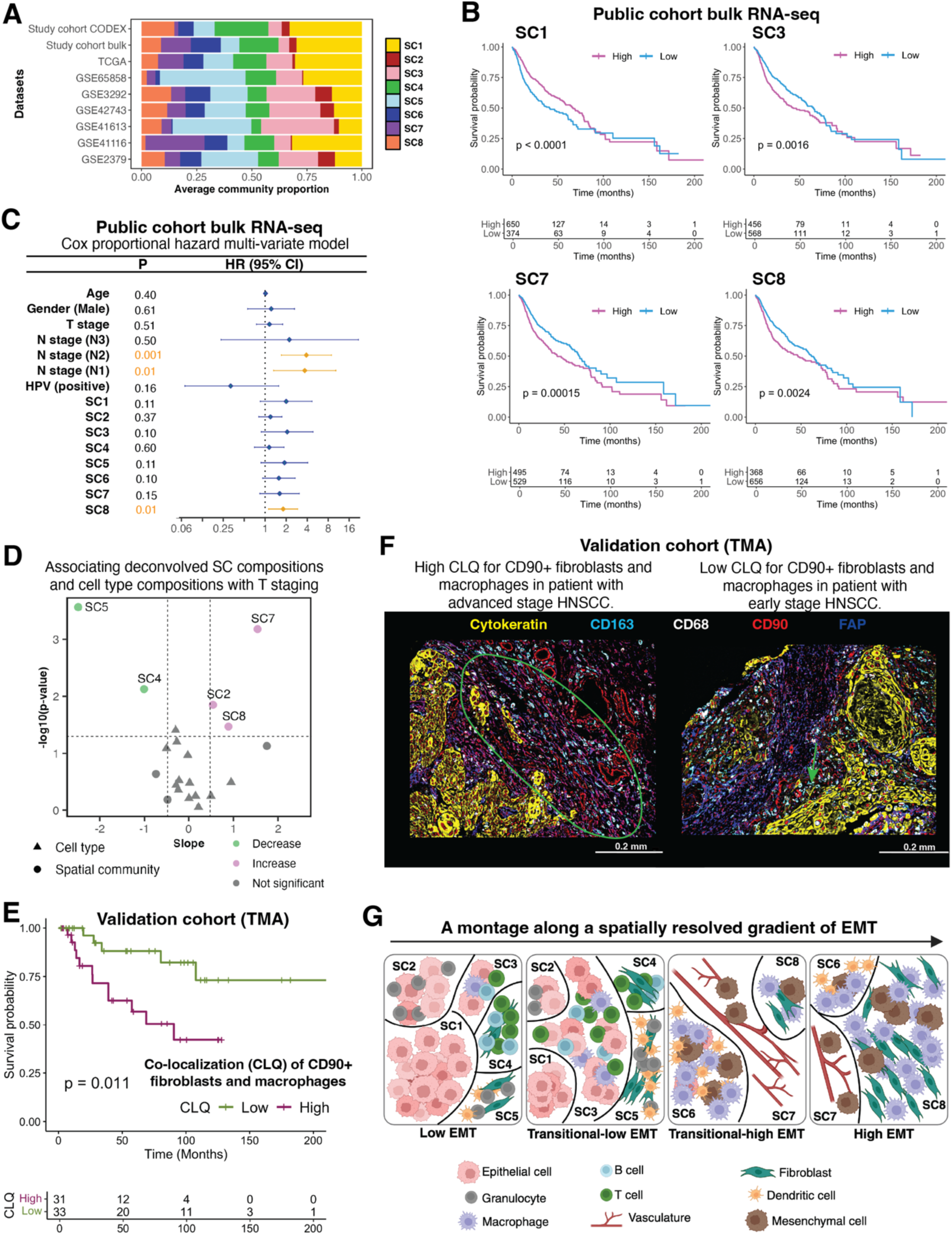
MONTAGE signature matrix used to deconvolve public bulk RNA-seq datasets into spatial community compositions for clinical relevance and validation. **(A)** Barplot showing high concordance of the average spatial community (SC) compositions of the CODEX data in the study cohort and average deconvolved SC compositions across multiples bulk RNA-seq datasets. **(B)** Kaplan-Meier curve survival analysis identified multiple prognostic MONTAGE deconvolved public bulk RNA-seq SCs. **(C)** Cox proportional hazard multi-variate survival analysis of deconvolved SC compositions and clinical factors. HR: hazard ratio. P: p-value. CI: confidence interval. **(D)** Comparison between deconvolved SC compositions and deconvolved cell type compositions association with T staging. **(E)** Kaplan-Meier survival analysis of co-localization between CD90+ fibroblast cells and macrophages measured by co-location quotient (CLQ) on the TMA validation cohort. **(F)** Representative CODEX images of TMA cores of the validation cohort from a patient of advanced stage HNSCC and poor survival outcome with high CLQ of CD90+ fibroblast cells and macrophages (left), and a patient of early stage HNSCC and good survival outcome with low CLQ of CD90+fibroblast cells and macrophages (right). **(G)** Graphical illustration of SC reorganization along spatially resolved gradient of EMT in HNSCC, representing a montage along spatially resolved EMT gradient created through the MONTAGE framework.

We then performed Kaplan-Meier (KM) survival analysis for each individual SC composition across all clinical samples in the large (N > 1,000) HNSCC public RNA-seq dataset (**Fig 6B, Supplementary Fig S7B**). HNSCC patients with higher compositions of tumor cell-enriched SCs (e.g., SC1) were associated with a better survival outcome, whereas patients with higher proportions of vasculature-enriched SCs (e.g., SC7) and stromal-macrophage-enriched SCs (e.g., SC8) were associated with worse survival outcomes. We also leveraged bulk RNA-seq samples from the public HNSCC study cohort for which clinical factors including age, gender, staging, and human papillomavirus (HPV) status were available (N=146), to perform a survival analysis using a multivariate Cox proportional hazards model ^26^ with clinical co-variates and SC compositions. SC8 was the only SC significantly associated with worse survival outcomes independent of known clinical factors (**Fig 6C**). Collectively, these findings indicate that changes in the TME are associated with cancer progression. Of note, we also analyzed TCGA HNSCC mutation data and found that common driver mutations (i.e., genes with a mutation frequency higher than 15%) occurred independently of SC abundance and distribution (**Supplementary Fig S7C**); these findings suggest that factors other than known driver mutations contribute to TME changes associated with cancer progression.

To further dissect the MONTAGE-identified association of fibroblasts and macrophage (SC8) interactions with HNSCC progression, we analyzed both protein expression profiles in different cell types in the SCs from the imaging data of the study cohort, and identified that malignant cells expressed high vimentin and fibroblast cells expressed high CD90 in SC8 (**Supplementary Fig S7D-E**). CellChat ^32^ analysis of the scRNA-seq data in the study cohort revealed that CD90 expression on fibroblasts and integrin expression on macrophages were uniquely mediating potential crosstalk between fibroblasts and macrophages (**Supplementary Fig S8A-C**). Public Visium data also showed consistent CellChat results of this potential fibroblast and macrophage crosstalk in the EMT high regions (**Supplementary Fig S8D-F**).

### MONTAGE provides prognostic information not available from cell-type-only spatial analyses

We next compared SC deconvolution by MONTAGE with traditional cell-type deconvolution. We applied CIBERSORTx ^33^ by first building a cell-type gene signature matrix using only scRNA-seq data from our HNSCC study cohort and then deconvolved the HNSCC public bulk RNA-seq datasets into cell-type compositions. We associated these deconvolved cell type compositions and MONTAGE deconvolved SC compositions with clinical T stage scores that indicate the size of a primary tumor (see **Methods** section). Two SC community compositions (SC4, SC5) were significantly associated with smaller tumor sizes, and three others (SC2, SC7, SC8) were significantly associated with larger tumor sizes. However, the cell-type compositions had no statistically significant association with tumor size (**Fig 6D**), further emphasizing the importance of cell organization (not only size or number) in tumor progression and highlighting the added value of dynamic monitoring of spatial-omics data using MONTAGE. In addition, we performed KM survival analysis with deconvolved cell-type compositions (**Supplementary Fig S9A**). Our data indicate that although deconvolved compositions of certain individual cell types were not prognostic for HNSCC, deconvolved SCs enriched with these cell types were prognostic, again highlighting the importance of cell-type spatial organization. Collectively, these analyses demonstrate the significance of MONTAGE for identifying spatial cellular organizations linked to disease progression and prognosis, which traditional cell-type deconvolution methods did not detect.

### Validation of MONTAGE-derived prognostic significance of fibroblast and macrophage enrichment on an independent cohort via a tissue microarray (TMA)

We leveraged our TMA of 68 HNSCC patients with its targeted DNA sequencing and available survival data ^34^ for validation purposes. We performed the same SC identification analysis on the TMA CODEX data and compared cell-type compositions in the TMA SCs with those in whole-slide CODEX SCs from the HNSCC study cohort (**Supplementary Fig S9B**). Five spatial communities (SC1, SC3, SC4, SC7, SC8) were present in the TMA cores, but three (SC2, SC5, SC6) could not be identified, potentially due to smaller tissue coverage of the TMA cores, which appear to lack certain immune- and stroma-enriched regions. The absence of these SCs in the TMA prevented us from estimating SC compositions consistent with the whole-slide imaging analyses. However, because the CODEX imaging panel for the TMA included CD90 (**Supplementary Table S4**), we leveraged the TMA as an independent validation of the prognosis significance of CD90-expressing fibroblasts and macrophages in SC8. For this analysis, we used our previously published geospatial statistical strategy co-location quotient (CLQ) ^35^ to quantify CD90+fibroblast and macrophage spatial co-localization within each TMA core and associated the CLQ values with survival outcomes. Patients with higher CD90+ fibroblast and macrophage CLQs had worse survival outcomes (**Fig 6E**). With a multivariate Cox proportional hazard model, we showed that CD90+fibroblast and macrophage co-localization was significantly prognostic, independent of major clinical factors in our validation cohort (**Supplementary Fig S9C-E**), consistent with the results from the MONTAGE-based deconvolved public bulk HNSCC RNA-seq datasets. Representative images of an advanced-stage patient and an early-stage patient from the validation cohort are shown in **Fig 6F**, illustrating the different co-localization patterns of CD90+fibroblasts and macrophages and further demonstrating the ability of MONTAGE to determine dynamic relationships between SC compositions and functional changes in the TME (**Fig 6G**).

## Discussion

MONTAGE is a computational framework that identifies biologically and clinically significant TME SCs by leveraging an integrated tissue atlas generated from single-cell spatial proteomics on whole-slide tissue sections and single-cell RNA sequencing (scRNA-seq). MONTAGE’s spatial community (SC) gene signature matrix reconstructs a tissue map, assigning each cell to its SC and functional enrichment. MONTAGE then applies local Moran’s I spatial autocorrelation to identify regions of distinct functional enrichment that accompany increased activity of biological processes that contribute to cancer progression, immune response, or drug resistance. MONTAGE thus creates sequences of SC compositions (“montages”) along spatially defined transitions in which functional pathways shift. In addition, the MONTAGE signature matrix enables deconvolution of large external non-single-cell data cohorts (e.g., TCGA), augmenting traditional cell-type-only deconvolution methods. Use of the MONTAGE signature matrix for both approaches can illuminate key potential interactions among cancer, stromal, and immune cells – and revealing their functional contributions to disease progression.

Applying MONTAGE to analysis of HNSCC advances understanding of the sixth most common cancer worldwide with around 890,000 new incidences in 2022 ^36,37^, and suggests how different SCs contribute to HNSCC progression. MONTAGE identified decreasing levels of tumor cell-, granulocyte- and lymphocyte-enriched SCs and increasing levels of stromal- and innate immune cell-enriched SCs along the EMT gradient. These findings demonstrate how the TME reorganizes its cellular composition as tumors progress toward a more mesenchymal, increasingly malignant state. Of note, MONTAGE identified a fibroblast- and macrophage-enriched community (SC8) that dominates regions of high-EMT enrichment. This finding is consistent with known pro-tumorigenic roles of fibroblasts and macrophages in promoting invasion and metastasis ^27^, concepts that emerged independently from our study cohort and validated in an independent spatial transcriptomics cohort.

MONTAGE was also instrumental for establishing the clinical relevance of SCs by deconvolving bulk RNA-seq and spot-based spatial transcriptomics HNSCC datasets using its signature matrix. Applying the MONTAGE gene signature matrix derived from our study cohort to deconvolve a large HNSCC patient cohort (N > 1000), we found SC8 proportions strongly correlated with worse survival outcomes, independent of common clinical covariates. These findings were confirmed by analyzing an independent TMA of 68 HNSCC patients, in which high co-localization of CD90+ fibroblasts and macrophages, the dominant cell types of SC8, were associated with poor survival. Our analysis also demonstrated that common driver mutations were independent of SC compositions. These results thus underscore the importance of searching beyond mutation-based prognostic markers and highlight the critical role of TME architecture in cancer progression.

Because MONTAGE employs a SC signature matrix that augments traditional cell type-only signatures, the approach is particularly beneficial for analyzing tumors in which interactions among diverse cell types are crucial drivers of pathology. While MONTAGE can be applied to standard TMA cores, we demonstrate the value of MONTAGE for analysis of whole-slide images. Of note, our data show that several stromal- and immune-enriched HNSCC SCs were not captured in our TMA-based HNSCC cohort, underscoring the value of analyzing whole-slide samples that contain non-cancer cells to fully understand the role of the TME in cancer progression. MONTAGE facilitates spatial-gradient functional analysis, enabling visualization of how different SCs expand or contract in regions of varying functional activity and thus offering a dynamic rather than static view of the TME. MONTAGE is scalable for analysis of bulk RNA-seq datasets, effectively addressing current sample-size limitations inherent in single-cell spatial imaging. By enabling SC deconvolution of thousands of bulk RNA-seq samples, MONTAGE allows for the association of SC compositions with critical clinical endpoints such as patient survival, tumor stage, and other relevant metrics.

### Limitations of the study

Although MONTAGE advances analysis of the TME in many ways, we acknowledge several limitations. As presented, MONTAGE is primarily designed to analyze deeply profiled cohorts in which cost and technical constraints often limit whole-slide single-cell studies. MONTAGE leverages the integration of moderately multiplexed images and scRNA-seq data to build an SC gene signature matrix; however, formation of this signature matrix depends on adequately powered scRNA-seq data. Uneven cell coverage in either scRNA-seq or imaging datasets could limit capturing the full diversity of SCs. To address these limitations, the MONTAGE framework can be extended for use with more comprehensively multiplexed single-cell imaging approaches such as Xenium or CosMx ^38^. MONTAGE can also be expanded to incorporate additional spatial assays (e.g., spatial metabolomics), because integrating higher-dimensional data might uncover additional functional gradients in the TME important for cancer progression or functionally specialized cellular niches.

In summary, MONTAGE provides a powerful framework to analyze cellular communities, link these communities to functional processes, and demonstrate their clinical significance. By prioritizing whole-slide tissue analysis, MONTAGE captures the complexity of tissue microenvironments more comprehensively than cell type-only analyses, making it broadly applicable to understanding normal and pathological processes in any tissue. Through its integration of single-cell imaging, scRNA-seq, functional gradient mapping, and large-scale deconvolution, MONTAGE reveals clinically relevant biological functions underlying the intricate spatial organization of the TME and various other tissues.

#### STAR Methods

### Creating the MONTAGE spatial community (SC) gene signature matrix

The MONTAGE framework begins with creating a SC gene signature matrix by integrating scRNA-seq and single-cell spatial proteomics data. MONTAGE requires two input matrices, matrix ***G*** derived from scRNA-seq and matrix ***S*** derived from multiplexed imaging:

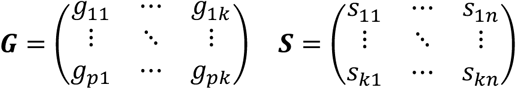

Where *p* is the number of cell type gene markers, *k* is the number of cell types, and *n* is the number of spatial communities. Matrix ***G*** is a *p* × *k* matrix that contains average gene expression *g* for each gene marker for each cell type. Matrix ***S*** is a *k* × *n* matrix that has the proportion *s* of each cell type in each spatial community. Descriptions of how to construct matrices ***G*** and ***S*** using our study cohort are described below. MONTAGE generates a spatial community gene signature matrix ***M*** by multiplication of matrices ***G*** and ***S***, which encodes each spatial community by weighting cell type marker gene expressions with corresponding cell type proportions in each spatial community.

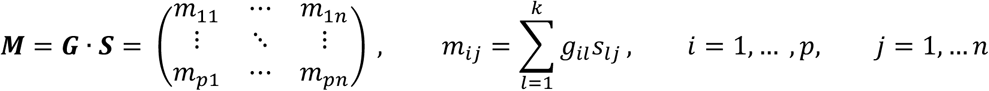

Please note that, the matrix ***G*** and matrix ***S*** can also be generated from spatial transcriptomics platforms with single-cell resolution and high transcriptomic coverage, to create MONTAGE SC gene signature matrix ***M*** using the same framework.

### Building the HNSCC study cohort

All patients included in our HNSCC study cohort from Stanford Hospital gave consent to participate in this study without compensation following Institutional Review Board (IRB) approval (IRB protocol no.11402). The patient information is summarized in Table S1. Fresh whole-slide tissue samples of HNSCC primary tumors and metastatic lymph nodes were included. Protocols for tissue preparation, single-cell RNA sequencing, CODEX imaging of the primary tumor sample data were published in a previous study ^35^. Metastatic lymph node data presented in this current study were generated concurrently with the primary tumor data following the same protocols. The bulk RNA-seq data were newly generated for this study. Bulk RNA was processed from a small slice of OCT tissue proximal to the slice used for CODEX data generation and sent to Medgenome for RNA extraction and sequencing. RNA extraction was performed using the Takara SMARTer stranded total RNA-seq kit v2, and library preparations were made using the Takara SMART-Seq v4 ultra low input RNA kit. Quality controls were run on RNA and library preparations using the Qubit fluorometric quantitation tape station bioanalyzer. Bulk sequencing was done on a NovaSeq (PE100/150) with 40M total reads/sample.

We created an HNSCC single-cell atlas of scRNA-seq, CITE-seq and CODEX data. Please note that only scRNA-seq and spatial proteomics data are required in the MONTAGE framework. Single-cell RNA-seq and CITE-seq data were processed using Cell Ranger v3.0.2 (10x Genomics) for initial alignment and quantification. Three different CITE-seq antibody panels were used across samples: a 36-plex panel (scc7239, scc7239B), a 44-plex panel (scc7268, scc7267B), and a 49-plex panel (scc7275, scc7276B) (Table S5). Further analysis was performed using Seurat v3.1.1 in R. Quality control filtering was applied to remove low-quality cells and likely doublets based on number of genes detected, number of UMIs, and percentage of mitochondrial reads (minimum 100 genes detected; maximum 10% mitochondrial reads, 5,000 genes detected, and 50,000 RNA counts). Data were normalized using SCTransform ^39^ and batch effects were corrected using Seurat’s integration method ^8^. Dimensionality reduction was performed using PCA followed by UMAP for visualization. CITE-seq log counts were integrated to remove batch effects and missing markers were imputed using Seurat’s anchor-based cross-modality transfer learning approach based on canonical correlation analysis (CCA). CITE-seq data were examined as histograms, as correlation to RNA expression for the corresponding gene, and in the UMAP space throughout data normalization, integration, and imputation to assess results in each step. Unsupervised clustering was carried out using Seurat’s Louvain algorithm. Cell types were annotated, and doublets were removed, based on expression of known marker genes and proteins. CODEX data were preprocessed to remove background and normalize marker expression levels. Cells were segmented based on nuclear staining and filtered to prepare a high-quality CODEX data subset for integration with single-cell sequencing data (minimum 2 proteins detected, 10,000 count, 10 sum of normalized values, and homogeneity of 3; maximum 30 proteins detected, 250,000 count, 70 sum of normalized values, 10,000 DNA score, and 5,000 cell size). Protein expression values were transformed using arcsinh, with scale factor for each protein determined based on histogram expression profiles, and integrated to remove batch effects. Dimensionality reduction was performed using PCA followed by UMAP for visualization. CODEX data were examined as histograms and in the UMAP space throughout data normalization and integration. To identify similar cell types in the sequencing and imaging modalities, we identified anchors between the two datasets using Seurat’s CCA. To do this, we used all CITE-seq surface proteins that overlapped with the CODEX panel, while using RNA expression of the remaining CODEX markers, including expression of all intracellular antigens that could not be measured by CITE-seq. This allowed for a reference-based label transfer approach to propagate cell type annotations from the scRNA-seq/CITE-seq data to the CODEX data. Please note that in the atlas, the scRNA-seq and CODEX data had correspondent cell types except granulocytes, which were only present in the CODEX data. CODEX and scRNA-seq data exhibited different cell type compositions across samples (Fig 2A-B). The differences may be due to cell dissociation process which could over-enrich lymphocyte populations in the scRNA-seq data ^40^.

To create matrix ***G*** from scRNA-seq data for building MONTAGE spatial community gene signature matrix, significant differentially expressed gene (DEGs) between each scRNA-seq cluster and all other clusters were identified using the Seurat FindMarkers function. Top one hundred genes with the highest fold changes were selected in clusters if more than one hundred DEGs were identified. An average expression for each gene was calculated across the cells for each cell type to construct matrix ***G***. For granulocyte population which was not present in the scRNA-seq data, gene expressions for granulocytes in the LM22 cell signature matrix in CIBERSORT ^11^ were scaled and included in matrix ***G***.

### Spatial community identification in single-cell resolution spatial proteomics data

We adapted a common strategy used in the current literature ^19,20^ for spatial community identification in spatial proteomics data. Briefly, for a given cell, we defined a window by selecting the ten nearest-neighboring cells around that cell and calculated the cell type proportions in each window to create vectors of cell type compositions. We next performed k-means clustering on the cell type composition vectors across the ten tissue samples. The cell type composition vectors were initially over-clustered to 30 clusters. We next mapped the clusters back onto the tissue samples for manual assessment and merged the clusters into 14 spatial neighborhoods quantifying densities of cell types in local proximity. To obtain the spatial communities, we next calculated the cell type compositions in each neighborhood and performed k-means clustering on the 14 spatial neighborhood cell type compositions. To define the optimal number of spatial communities, we constructed spatial community gene signature matrices based on community numbers from 5 to 13, and deconvolved the spatial community proportions of the bulk RNA-seq samples generated from adjacent sections with the CODEX data in the study cohort. We calculated the cosine similarity scores between the deconvolved SC proportions and the SC proportions in the CODEX data which we considered as ground truth. The number of SCs was decided by the best similarity score (SC=8) obtained on the paired bulk RNA-seq data in the study cohort (Supplementary Fig S3C). Each SC was annotated with cell type enrichment based on the cellular compositions in that community.

Matrix ***S*** was constructed with cell type proportions for each cell type in each spatial community. To obtain the sample comparison of the deconvolved SC proportions from the bulk RNA-seq and the ground truth SC proportions, we calculated the Cosine similarity score and categorized the proportions to calculate a Rand Index for each sample.

### Creating a montage of spatial community composition changes along spatially resolved gradient of differential functional enrichment

We applied GSVA ^21^ to calculate gene set enrichment scores for the gene signatures in each SC in the MONTAGE HNSCC signature matrix. For illustration purpose, we focused on the 50 genesets included in the MSigDB Hallmark genesets ^22^ representing well-defined biological states or processes. We obtained gene set enrichment scores for each hallmark of each SC (Fig 3D). Each individual cell in the CODEX data was assigned its corresponding SC enrichment scores. MONTAGE can superimpose each enrichment scores for individual cells onto the CODEX whole-slide images, which allows displaying any specific gene functional enrichment in spatial context at single-cell resolution.

To create montages along spatially resolved gradients on the functional enrichment scores, we first calculated local Moran’s I ^24^ for each individual cell in the CODEX data using the R package “spdep”. Local Moran’s I is a geospatial statistical measure indicating the extent of significant spatial clustering of similar values around an observation. For a specific functional enrichment for each cell, the local Moran’s I was calculated based on scores of 100 nearest neighboring cells around that cell. After obtaining the local Moran’s I statistics for each cell, we can define four regions along a spatially resolved gradient of a functional enrichment, namely, region with negative enrichment scores and significant local Moran’s I, region with negative enrichment scores and non-significant local Moran’s I, region with positive enrichment scores and non-significant local Moran’s I and region with positive enrichment scores and significant local Moran’s I. In the example of functional enrichment for EMT, we identified four regions annotated as low EMT, transitional-low EMT, transitional-high EMT and high EMT regions (Fig 4A). The low EMT and high EMT regions had significant local Moran’s I with low and high EMT scores respectively, which indicated significant spatial clustering of low and high functional enrichment of EMT. The two transitional regions did not show significant local Moran’s I, suggesting spatially intermixing of EMT scores. Within each region, we calculated the average composition for each SC across the ten CODEX samples, which thereby provided quantitation on how the SC compositions progressed along the spatially resolved gradient of EMT in HNSCC whole-slide tissue. We also performed the same analysis on glycolysis to illustrate the capability of MONTAGE to study any biological function enrichment in tissue.

### Applying MONTAGE to deconvolve public Visium data

A public HNSCC Visium dataset (GSE208253) with 12 whole-slide samples was used to demonstrate MONTAGE’s application on spot-based spatial transcriptomics data. Seurat objects of processed transcriptomic expressions included in the public datasets were used for deconvolution and creating montages. We applied MONTAGE HNSCC signature matrix generated from our study cohort in the deconvolution using CIBERSORT ^11^. We first deconvolved the three main pathologist annotated regions included in this public dataset, namely, SCC tumor, lymphocyte negative stroma and lymphocyte positive stroma. We showed that the deconvolved SC compositions agreed with pathologist annotations. We next used Seurat to cluster the RNA expressions with FindClusters function and identified spatial clusters for each sample, and we applied CIBERSORT with MONTAGE HNSCC signature matrix to deconvolve each spatial clusters. While it is possible to deconvolve each spot in the Visium with MONTAGE HNSCC spatial community gene signature matrix, we opted to perform deconvolution on spatial clusters to reduce computational complexity.

We calculated functional enrichment score for EMT of each spot using GSVA. We applied local Moran’s I to identify the four regions along the spatially resolved gradient of EMT enrichment scores. Given that each spot typically contains 10 to 20 cells in Visium data, we defined neighborhoods using 5 nearest spots in the local Moran’s I calculation, which was comparable to the neighborhood of 100 nearest cells defined in the single-cell resolution CODEX data from our study cohort. We used the same criteria of combining EMT enrichment scores and local Moran’s I significances to categorize the Visium spots into low EMT, transitional-low EMT, transitional-high EMT and high EMT regions. Within each EMT region, we calculated the average composition for each SC across the 12 Visium samples, resulting in similar quantifications of SC composition progression along the spatially resolved gradient of EMT as observed in the CODEX data in our study cohort.

SC proportions between patients with (N+) and without (N0) lymph node metastasis were compared using T-test. DEG analysis between EMT high and EMT low regions were performed using a consensus strategy ^41^. Briefly, gene expressions between high and low EMT regions were compared using Wilcoxon test with Bonferroni correction for multiple comparisons. Genes considered to be differentially expressed had adjusted p-values smaller than 0.05 and log fold changes higher than 0.2 in at least 6 out of the 12 samples. CellChat v2 ^42^ for spatial transcriptomics data was applied to identify ligand-receptor pairs in the four EMT regions.

### Applying MONTAGE to deconvolve public bulk transcriptomics data

To quantify transcriptomic expression of the bulk RNA-seq data from our HNSCC study cohort, we used Kallisto v0.46.1 ^43^ to align reads to the GENCODE human transcripts v43. Transcript-level expressions were summarized to gene level by TPM values using tximport ^44^. The processed gene level expressions of other public HNSCC bulk RNA-seq datasets were downloaded from either GEO or TCGA portal (Data availability). We applied CIBERSORT to deconvolve each bulk RNA-seq datasets into their SC compositions using the MONTAGE signature matrix generated from our HNSCC study cohort. Average spatial community proportions for each dataset were compared with the CODEX SC compositions in our study cohort using Cosine similarity scores. The Kaplan-Meier (KM) survival analysis was performed on all public bulk RNA-seq samples (n=1028) using normalized SC proportions by survfit function in R survival package. Median values were used to define the high and low spatial community composition groups. The Cox proportional hazard multi-variate survival analysis was performed by coxph function in R survival package using samples from datasets where survival outcome, HPV status, gender, staging and age information were available (TCGA, GSE3292, GSE41116). The SC compositions were included in the Cox model as continuous variables. TCGA mutation data was used to assess the SC compositions with respect to mutation status. Chi-square test was applied on genes with mutation frequency higher than 15%.

To compare the traditional cell-type deconvolution and MONTAGE spatial community deconvolution, we used CIBERSORTx ^33^ to derive a HNSCC cell-type gene signature matrix from the scRNA-seq data in our HNSCC study cohort. We applied CIBERSORT to deconvolve the bulk RNA-seq datasets into cell type compositions with this HNSCC cell-type signature matrix. To associate the deconvolved cell type compositions and SC compositions with staging, we leveraged the samples from datasets with T stage annotations (TCGA, GSE3292, GSE41116). We performed ANOVA tests of cell type proportions and SC proportions against T stages to assess significant association. We used coefficients from linear regression to define the association directions. We performed the same KM curve survival analysis using the deconvolved cell type compositions on all the bulk transcriptomic samples with median values to define high and low composition groups.

### THY1 (CD90) and integrin crosstalk analysis

Segmented expressions of CD90, FAP, Vimentin and Cytokeratin protein staining from the CODEX data in our study cohort were compared across spatial communities. Each expression was arcsinh transformed. The expressions of CD90 and FAP of fibroblast cells in spatial communities SC2-7 were normalized against the expressions of CD90 and FAP of fibroblast cells in SC1 respectively to obtain log transformed ratio for comparison. Similarly, the expressions of Vimentin and Cytokeratin of malignant cells in spatial communities SC2-7 were normalized against the expressions of Vimentin and Cytokeratin of malignant cells in SC1 respectively to obtain log transformed ratio for comparison. CellChat ^32^ was applied to the scRNA-seq data from our HNSCC study cohort, and signaling between fibroblast cells and macrophages was analyzed and visualized.

### Validation HNSCC tissue microarray cohort analysis

An HNSCC tissue microarray (TMA) of oral cavity, HPV negative patients was constructed in a previous published study ^34^. Primary tumor cores, adjacent benign mucosal/stromal areas and metastatic/benign lymph nodes were sectioned at 4 μm and mounted onto Vectabond™-treated glass coverslips (21 mm × 21 mm). Commercially available, purified, carrier-free anti-human antibodies (Table S4) were conjugated to maleimide-modified DNA oligonucleotides and titrated on the tissue of interest following previously published protocols ^45^. All the antibodies used were previously validated for use in CODEX multicycle imaging. The multicycle image acquisition was conducted using Akoya’s CODEX instrument connected to a Keyence BZ-X700 microscope configured with four fluorescent channels (DAPI, FITC, Cy3, Cy5) and a 20x objective. TMA spots were imaged using 1×1 tile and multiple z planes. CODEX imaging data was processed using a software tool called RAPID ^46^ (available at: https://github.com/nolanlab/RAPID), which included 3D GPU-based deconvolution, spatial drift correction, image stitching, and autofluorescence background reduction available at and segmented using a public CODEX image pipeline available at https://github.com/nolanlab/CODEX. Cell type identification was performed using CELESTA ^35^. We used the same spatial community identification strategy described above for whole-slide tissue to group the cells on all the TMA primary tumor center cores into eight clusters. We compared the cell type compositions in each cluster with the cell type compositions in the eight SCs identified from whole-slide tissue samples using Cosine similarity scores. To quantify the spatial co-localization of CD90+ fibroblasts and macrophages, we adapted a geospatial statistical measure CLQ (co-location quotient) ^35^ for each TMA core. CLQ metric defines the degree of one cell type co-locates with the nearest neighborhood of another cell type. We divided the patient samples of the TMA into high and low CLQ groups by using the median value of all the CLQ values of CD90+ fibroblasts and macrophages across all the samples. The Kaplan-Meier survival analysis was performed by survfit function in R survival package on the two groups. We compared the mutation status of genes with mutation frequency higher than 3% in the targeted DNA sequencing data of this TMA with chi-square test between high CLQ and low CLQ groups. Leveraging the patient clinical information of this TMA, we applied Cox proportional hazard multi-variate survival analysis with gender, staging, age, TP53 mutation and CLQ values of CD90+ fibroblasts and macrophages co-localization included as the covariates.

### Figure creation

Figures were plotted in R and graphical illustrations in Figure 1, Figure 5A, Figure 6G were created with BioRender.com.

### Data availability

Whole-slide HNSCC tissue CODEX imaging data and corresponding segmented data included in the study cohort is accessible on Synapse at: https://doi.org/10.7303/syn26242593. HNSCC scRNA-seq data included in our study cohort can be accessed at Gene Expression Omnibus (GEO) with accession number GSE140042. HNSCC Bulk RNA-seq data included in the study cohort is accessible on GEO with accession number GSE286234. The CODEX data included in the validation cohort of HNSCC tissue microarray is available upon request. The TCGA HNSCC data was generated by the TCGA Research Network https://www.cancer.gov/tcga. Other HNSCC datasets included in the public cohort can be accessed at: GSE65858 (https://www.ncbi.nlm.nih.gov/geo/query/acc.cgi?acc=GSE65858). GSE3292 (https://www.ncbi.nlm.nih.gov/geo/query/acc.cgi?acc=GSE3292). GSE42743 (https://www.ncbi.nlm.nih.gov/geo/query/acc.cgi?acc=GSE42743). GSE41613 (https://www.ncbi.nlm.nih.gov/geo/query/acc.cgi?acc=GSE41613). GSE41116 (https://www.ncbi.nlm.nih.gov/geo/query/acc.cgi?acc=GSE41116). GSE2379 (https://www.ncbi.nlm.nih.gov/geo/query/acc.cgi?acc=GSE2379). GSE208253 (https://www.ncbi.nlm.nih.gov/geo/query/acc.cgi?acc=GSE208253).

### Code availability

The MONTAGE R package will be released on GitHub https://github.com/plevritis-lab/MONTAGE with public access upon acceptance of the manuscript.

## Author contributions

W.Z. and S.K.P. conceived and designed the computational framework. W.Z. implemented the computational framework and performed the data analysis. Z.G. created the HNSCC single-cell atlas of the study cohort and curated the public bulk HNSCC RNA-seq datasets. M.A.B., G.L., J.W.H, and G.P.N. generated the CODEX data for the validation cohort of HNSCC tissue microarray (TMA) and performed imaging preprocessing and segmentation. R.H. and Q.-T.L. curated the clinical information for the TMA of the validation cohort. S.C. performed tissue sample collection and generating bulk RNA-seq data of the HNSCC study cohort. A.J.G. and J.B.S. supervised the tissue sample collection and data generation for the study cohort. C.S.K. supervised and built the TMA validation cohort and provided pathologic support. S.K.P. supervised the overall project and study. All authors contributed to and approved the manuscript.

## Acknowledgements

This work was supported by the National Institutes of Health (NIH), National Cancer Institute U54CA209971 and U54CA274511. Z.G. was supported by the Stanford Cancer Institute Fellowship, Parker Scholar and Parker Bridge Fellow awards from the Parker Institute for Cancer Immunotherapy, and the NIH/NCI Pathway to Independence Award (1K99CA293149, 4R00CA293149). M.A.B. was supported by a Career Development award of the International Myeloma Society (IMS). J.W.H. was supported by an NIH T32 Fellowship (T32CA196585) and an American Cancer Society—Roaring Fork Valley Postdoctoral Fellowship (PF-20-032-01-CSM). Q.-T.L. was supported by 1R01DE029672, P01CA257907 and R01DE030894. J.B.S. was supported by R35DE030054. G.P.N. was supported by the US NIH P01HL108797, U01AI101984, 5U54CA209971, 5U01AI140498, U54HG010426, U19AI100627, R01HL120724, R01HL128173, 5P01AI131374, UH3DK114937, U19AI135976, U2CCA233195, U19AI057229, and U2CCA233238 and the Rachford Carlotta A. Harris Endowed Chair. We are very grateful for the patients who consented to use their tissues for this project. The authors also appreciate the editorial contributions of Alison F. Davis, PhD.

