## Supplementary Figures S1-S9 and Supplementary Tables S1, S3 for "MONTAGE: A Computation Framework to Identify Spatially Resolved Functional Enrichment Gradients in the Tissue Microenvironment via Spatial Communities"

### Supplementary Information

Supplementary Figures S1-S9 and Supplementary Tables S1, S3.
Supplementary Table S2, S4, S5: Excel files containing additional data too large to fit.

### Supplementary Figures

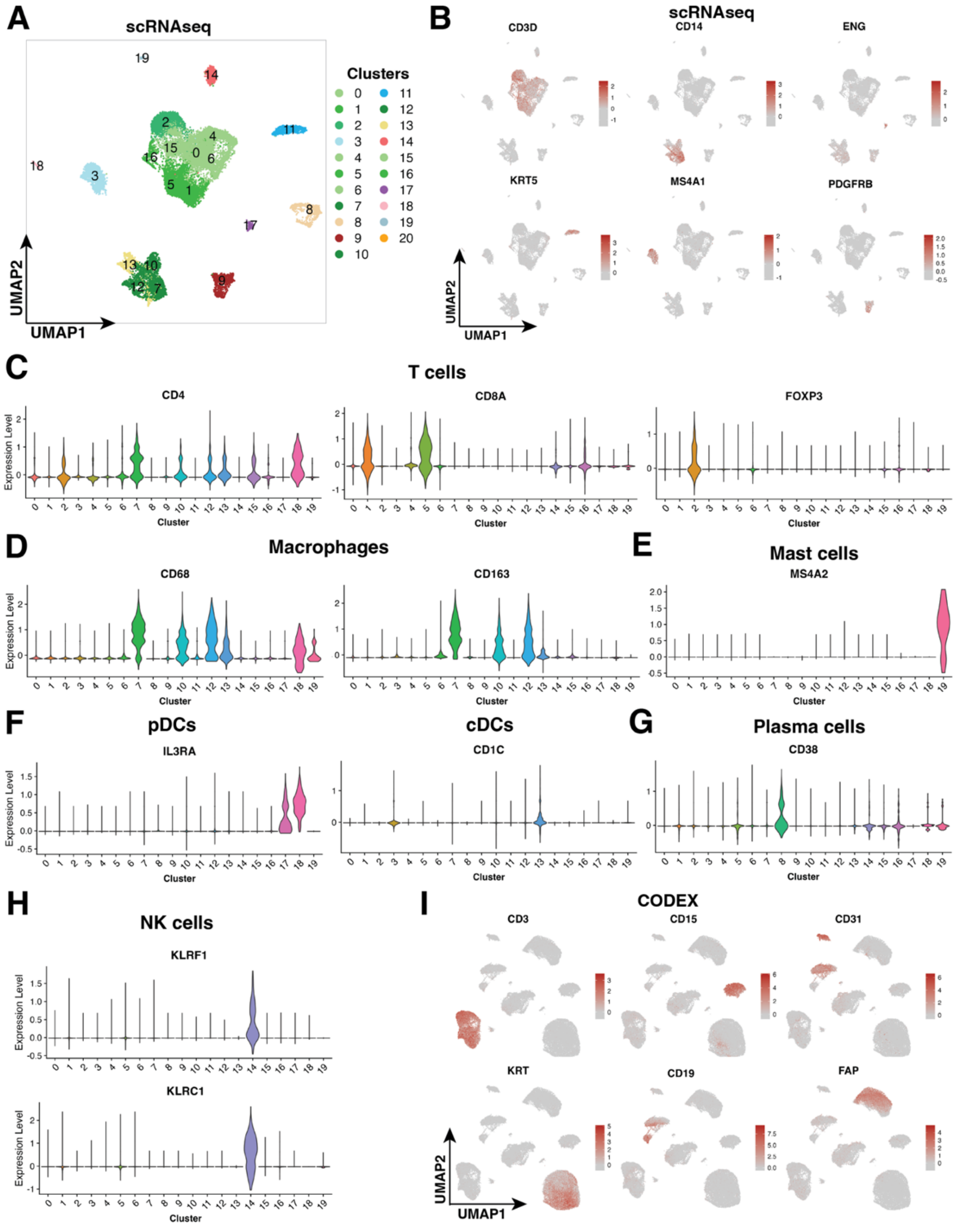

**Figure S1. Canonical marker expressions used for cell type assignments in the HNSCC study cohort.**
**(A)** UMAP of scRNA-seq clusters.
**(B)** UMAPs of major canonical markers of the scRNA-seq data.
**(C)-(H)** Canonical marker expressions for refined immune populations in each cluster.
**(I)** UMAPs of major canonical markers of the CODEX data.

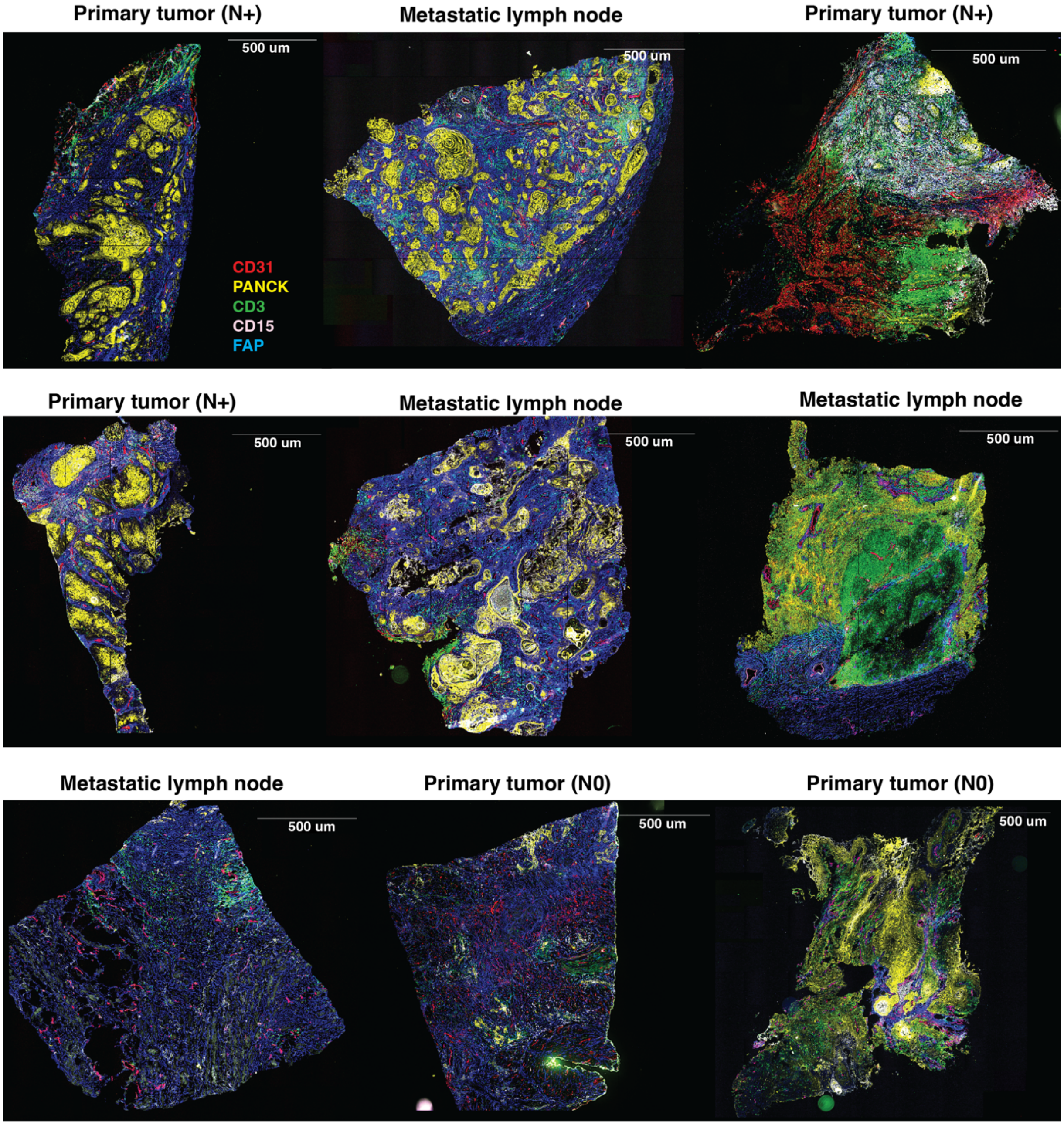


**Figure S2. Whole-slide CODEX images of representative canonical markers of samples in the HNSCC study cohort.**

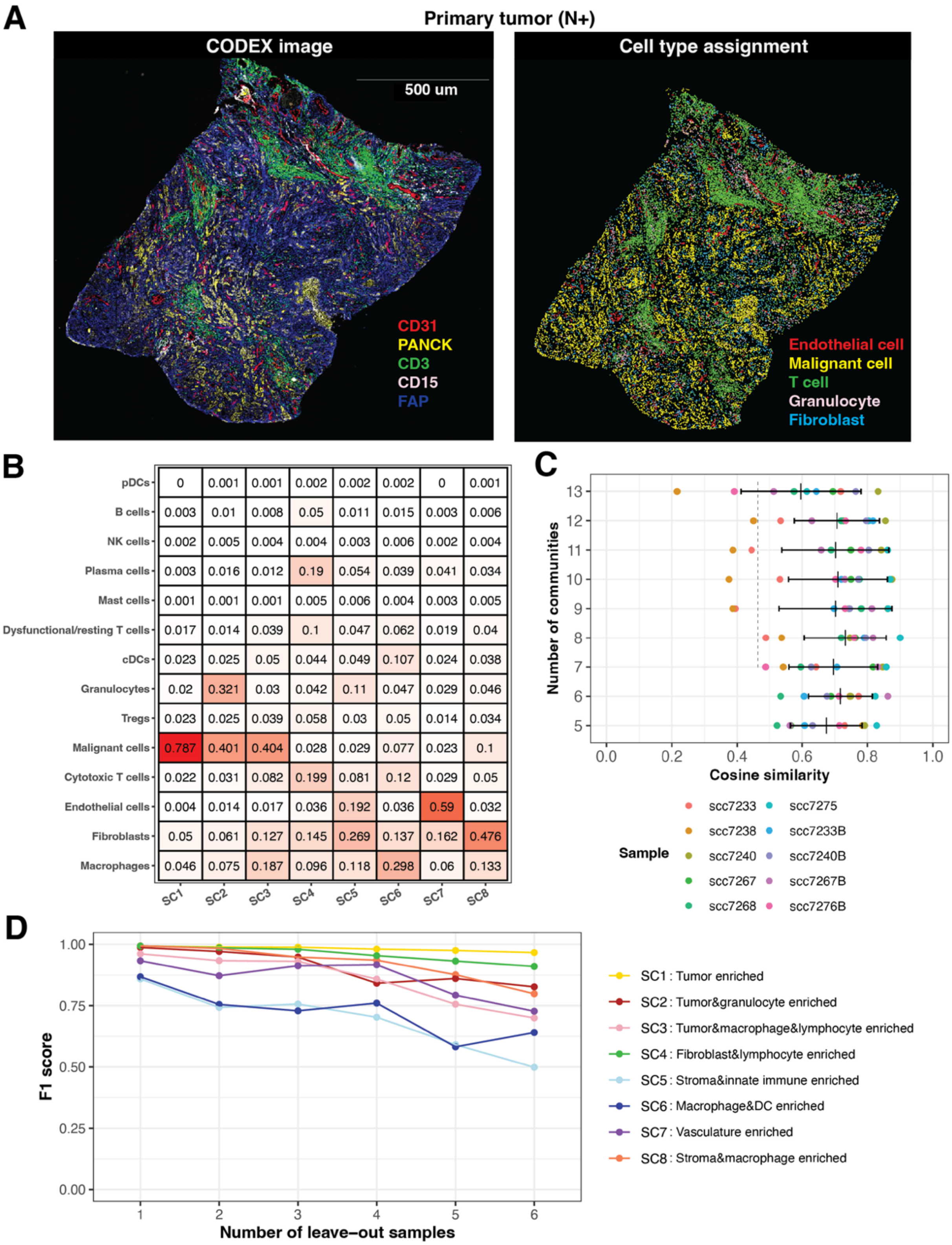
A
Primary tumor (N+)
CODEX image
500 um
CD31
PANCK
CD3
CD15
FAP
Cell type assignment
Endothelial cell
Malignant cell
T cell
Granulocyte
Fibroblast
B
pDCs 0 0.001 0.001 0.002 0.002 0.002 0 0.001
B cells 0.003 0.01 0.008 0.05 0.011 0.015 0.003 0.006
NK cells 0.002 0.005 0.004 0.004 0.003 0.006 0.002 0.004
Plasma cells 0.003 0.016 0.012 0.19 0.054 0.039 0.041 0.034
Mast cells 0.001 0.001 0.001 0.005 0.006 0.004 0.003 0.005
Dysfunctional/resting T cells 0.017 0.014 0.039 0.1 0.047 0.062 0.019 0.04
cDCs 0.023 0.025 0.05 0.044 0.049 0.107 0.024 0.038
Granulocytes 0.02 0.321 0.03 0.042 0.11 0.047 0.029 0.046
Tregs 0.023 0.025 0.039 0.058 0.03 0.05 0.014 0.034
Malignant cells 0.787 0.401 0.404 0.028 0.029 0.077 0.023 0.1
Cytotoxic T cells 0.022 0.031 0.082 0.199 0.081 0.12 0.029 0.05
Endothelial cells 0.004 0.014 0.017 0.036 0.192 0.036 0.59 0.032
Fibroblasts 0.05 0.061 0.127 0.145 0.269 0.137 0.162 0.476
Macrophages 0.046 0.075 0.187 0.096 0.118 0.298 0.06 0.133
SC1 SC2 SC3 SC4 SC5 SC6 SC7 SC8
C
Number of communities
13 12 11 10 9 8 7 6 5
0.0 0.2 0.4 0.6 0.8 1.0
Cosine similarity
Sample
scc7233
scc7238
scc7240
scc7267
scc7268
scc7275
scc7233B
scc7240B
scc7267B
scc7276B
D
F1 score
1.00 0.75 0.50 0.25 0.00
1 2 3 4 5 6
Number of leave-out samples
SC1 : Tumor enriched
SC2 : Tumor&granulocyte enriched
SC3 : Tumor&macrophage&lymphocyte enriched
SC4 : Fibroblast&lymphocyte enriched
SC5 : Stroma&innate immune enriched
SC6 : Macrophage&DC enriched
SC7 : Vasculature enriched
SC8 : Stroma&macrophage enriched

**Figure S3. Additional results on spatial community identification.**
**(A)** Left: A Whole-slide CODEX image of a primary tumor sample of a lymph node positive (N+) patient. Right: Selected cell types plotted on the same sample.
**(B)** Cell type proportions in each SC identified on the CODEX data.
**(C)**Testing different numbers of spatial communities by comparing similarity scores between the deconvolved SCs from bulk RNA-seq and SCs identified in the CODEX data (ground truth) in the study cohort.
**(D)** Leave-out sample analysis to test the robustness of the identified SCs.

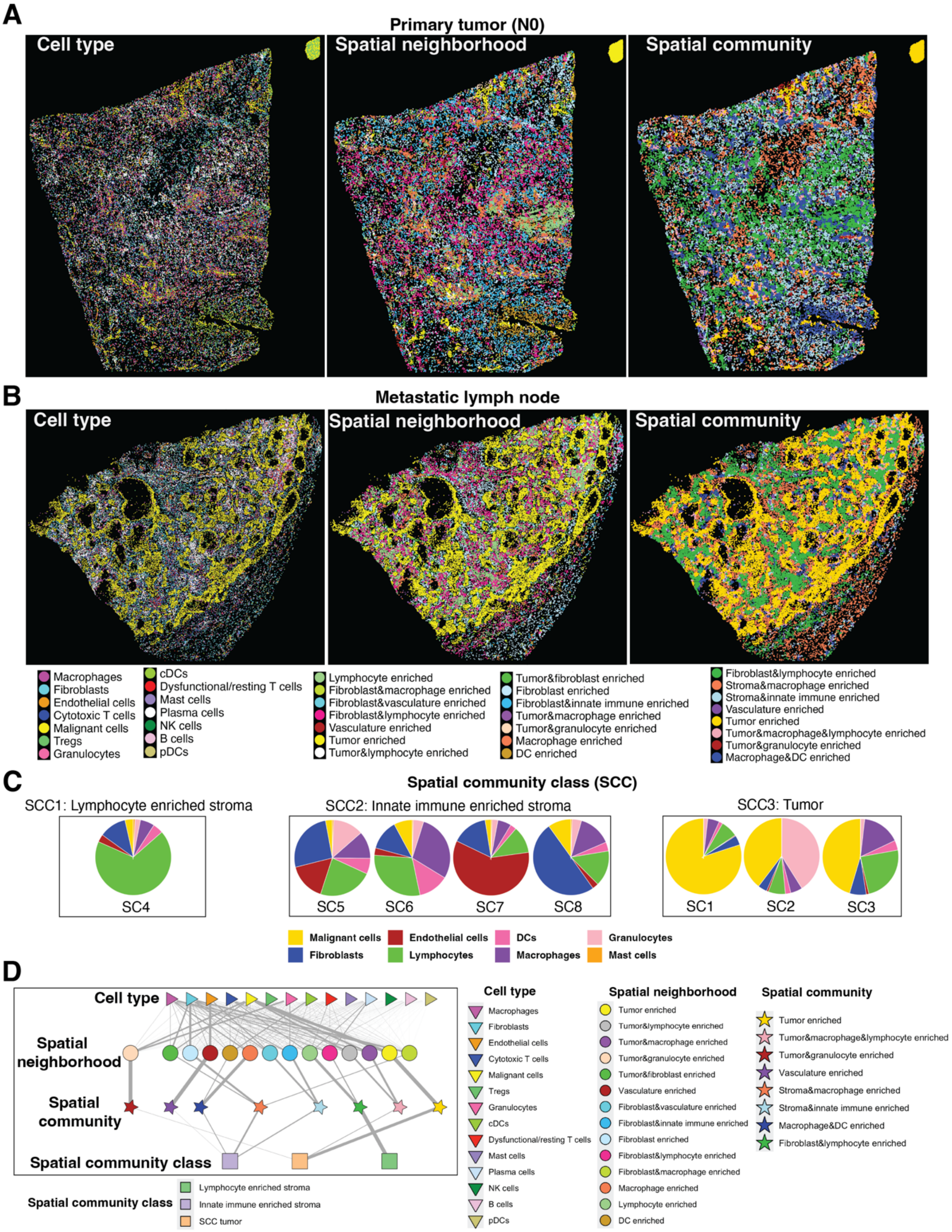

A
Primary tumor (N0)
Cell type
Spatial neighborhood
Spatial community
B
Metastatic lymph node
Cell type
Spatial neighborhood
Spatial community
Macrophages
Fibroblasts
Endothelial cells
Cytotoxic T cells
Malignant cells
Tregs
Granulocytes
cDCs
Dysfunctional/resting T cells
Mast cells
Plasma cells
NK cells
B cells
pDCs
Lymphocyte enriched
Fibroblast&macrophage enriched
Fibroblast&vasculature enriched
Fibroblast&lymphocyte enriched
Vasculature enriched
Tumor enriched
Tumor&lymphocyte enriched
Tumor&fibroblast enriched
Fibroblast enriched
Fibroblast&innate immune enriched
Tumor&macrophage enriched
Tumor&granulocyte enriched
Macrophage enriched
DC enriched
Fibroblast&lymphocyte enriched
Stroma&macrophage enriched
Stroma&innate immune enriched
Vasculature enriched
Tumor enriched
Tumor&macrophage&lymphocyte enriched
Tumor&granulocyte enriched
Macrophage&DC enriched
C
Spatial community class (SCC)
SCC1: Lymphocyte enriched stroma
SCC2: Innate immune enriched stroma
SCC3: Tumor
SC4
SC5
SC6
SC7
SC8
SC1
SC2
SC3
Malignant cells
Endothelial cells
DCs
Granulocytes
Fibroblasts
Lymphocytes
Macrophages
Mast cells
D
Cell type
Spatial neighborhood
Spatial community
Spatial community class
Cell type
Macrophages
Fibroblasts
Endothelial cells
Cytotoxic T cells
Malignant cells
Tregs
Granulocytes
cDCs
Dysfunctional/resting T cells
Mast cells
Plasma cells
NK cells
B cells
pDCs
Spatial neighborhood
Tumor enriched
Tumor&lymphocyte enriched
Tumor&macrophage enriched
Tumor&granulocyte enriched
Tumor&fibroblast enriched
Vasculature enriched
Fibroblast&vasculature enriched
Fibroblast&innate immune enriched
Fibroblast enriched
Fibroblast&lymphocyte enriched
Fibroblast&macrophage enriched
Macrophage enriched
Lymphocyte enriched
DC enriched
Spatial community
Tumor enriched
Tumor&macrophage&lymphocyte enriched
Tumor&granulocyte enriched
Vasculature enriched
Stroma&macrophage enriched
Stroma&innate immune enriched
Macrophage&DC enriched
Fibroblast&lymphocyte enriched
Spatial community class
Lymphocyte enriched stroma
Innate immune enriched stroma
SCC tumor

**Figure S4. Additional results of the spatial communities in the CODEX data from the study cohort.**
**(A)-(B)** Additional samples from a primary tumor sample of a lymph node negative (N0) patient and a metastatic lymph node sample showing the identified cell types, spatial neighborhoods and spatial communities (SCs).
**(C)** SCs combined into spatial community classes (SCCs) based on major cell type abundances.
**(D)** Hierarchical network showing the contribution from cell types to spatial neighborhoods, SCs and SCCs.

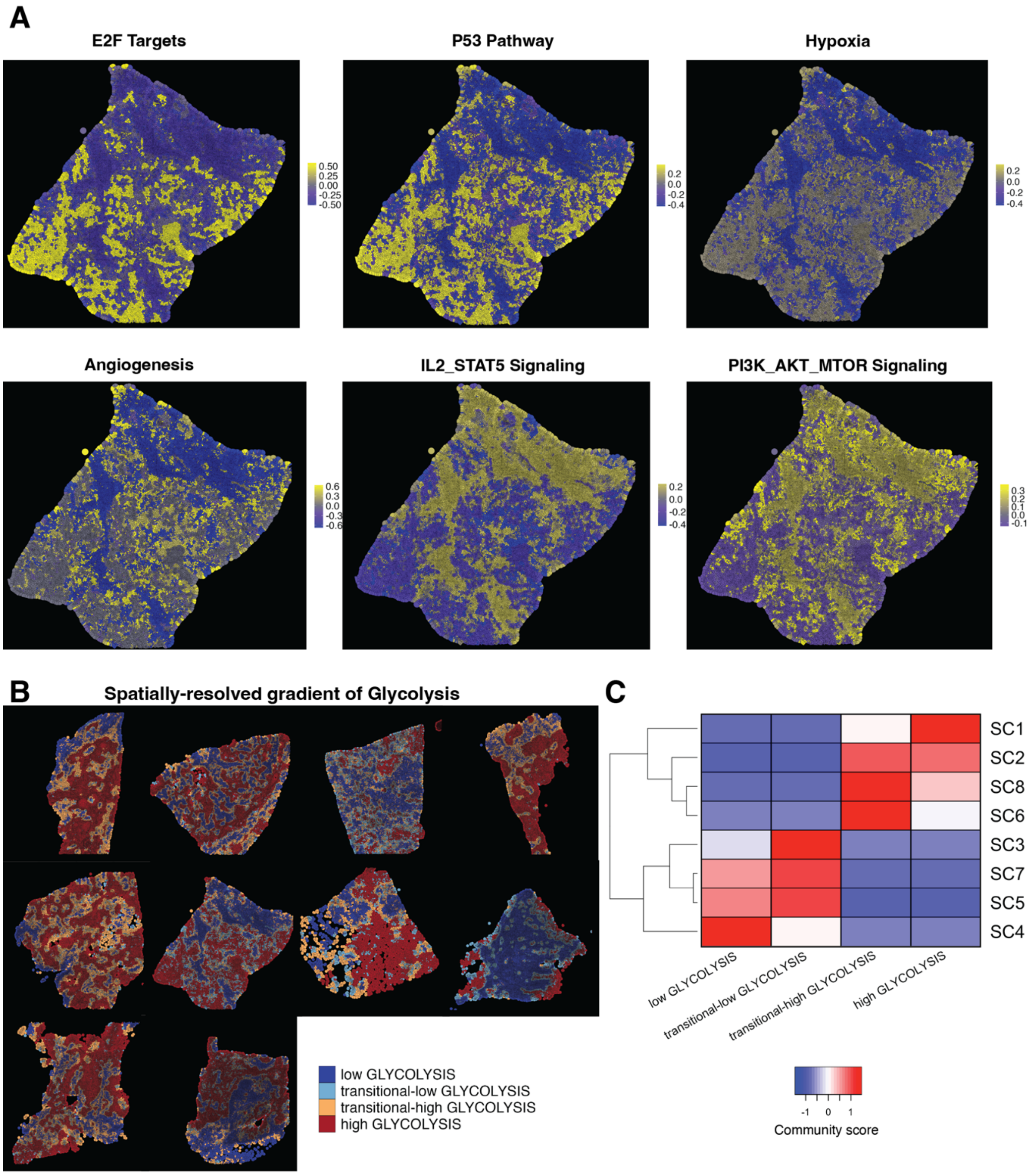


**Figure S5. Additional results of MONTAGE creating montages along spatially resolved gradient of functional enrichments.**

**(A)** Additional functional enrichment scores superimposed onto a representative whole-slide spatial proteomics image in the study cohort.

**(B)** Applying MONTAGE to identify spatially resolved gradient of increasing glycolysis enrichment scores.

**(C)** A heatmap shows normalized SC composition score progression along the spatially resolved gradient of glycolysis across 10 CODEX samples in the study cohort.

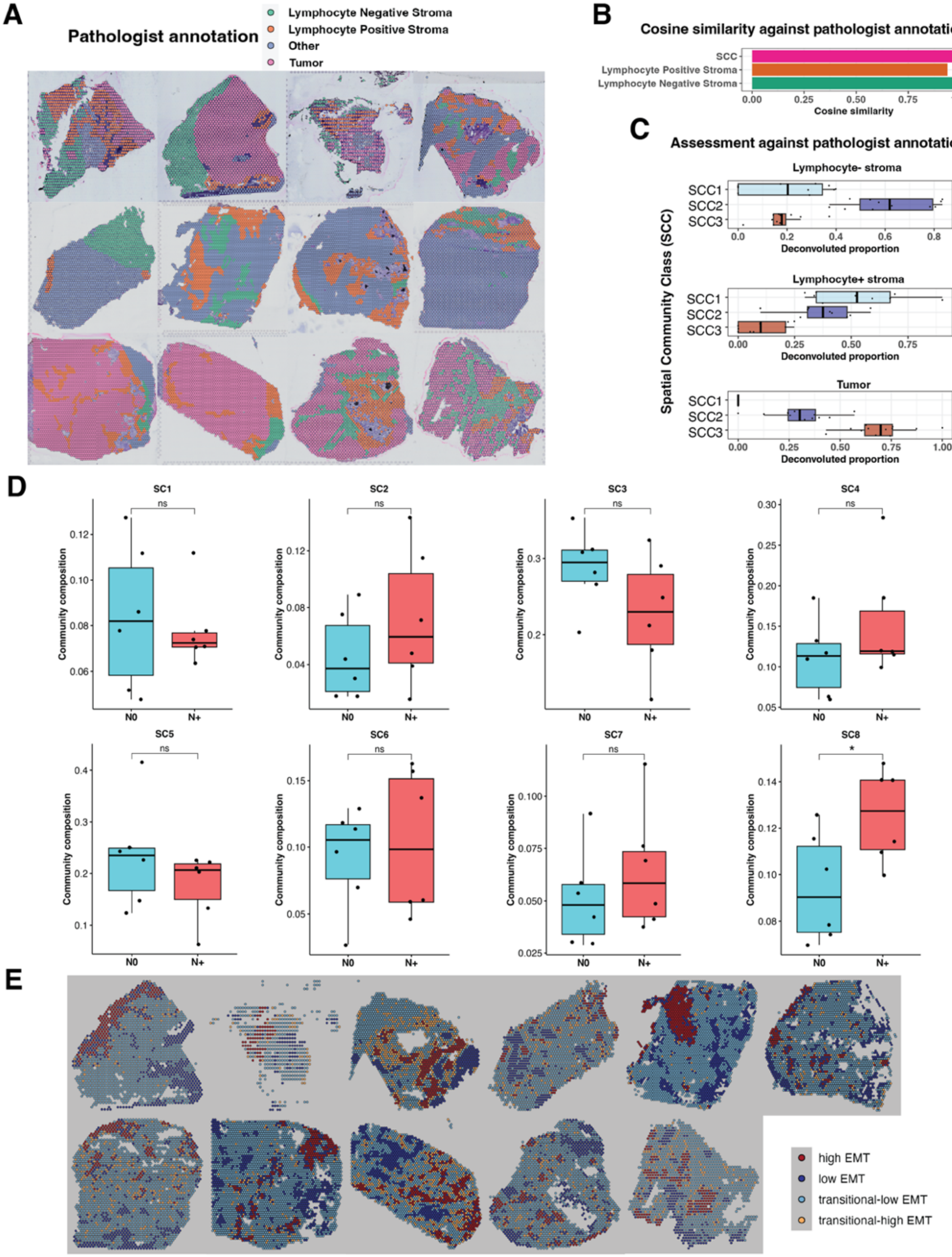

A
Pathologist annotation
Lymphocyte Negative Stroma
Lymphocyte Positive Stroma
Other
Tumor
B
Cosine similarity against pathologist annotation
SCC
Lymphocyte Positive Stroma
Lymphocyte Negative Stroma
Cosine similarity
C
Assessment against pathologist annotation
Spatial Community Class (SCC)
Lymphocyte- stroma
Lymphocyte+ stroma
Tumor
SCC1
SCC2
SCC3
Deconvoluted proportion
D
SC1
SC2
SC3
SC4
SC5
SC6
SC7
SC8
ns
*
Community composition
N0
N+
E
high EMT
low EMT
transitional-low EMT
transitional-high EMT

**Figure S6. Additional results of MONTAGE applied to deconvolve a public Visium dataset.**
**(A)** Pathologist annotations replotted on the 12 whole-slide tissue samples included in the public Visium data.
**(B)-(C)** Assessment of the deconvolved spatial community (SC) compositions against pathologist annotations.
**(D)** Barplot comparing SC compositions between lymph node negative (N0) and lymph node positive (N+) patients. Each condition has 6 samples. *: nominal p-value<0.05. ns: not significant.
**(E)** Additional samples of the four identified regions along spatially resolved gradient for increasing EMT enrichment in the public Visium data.

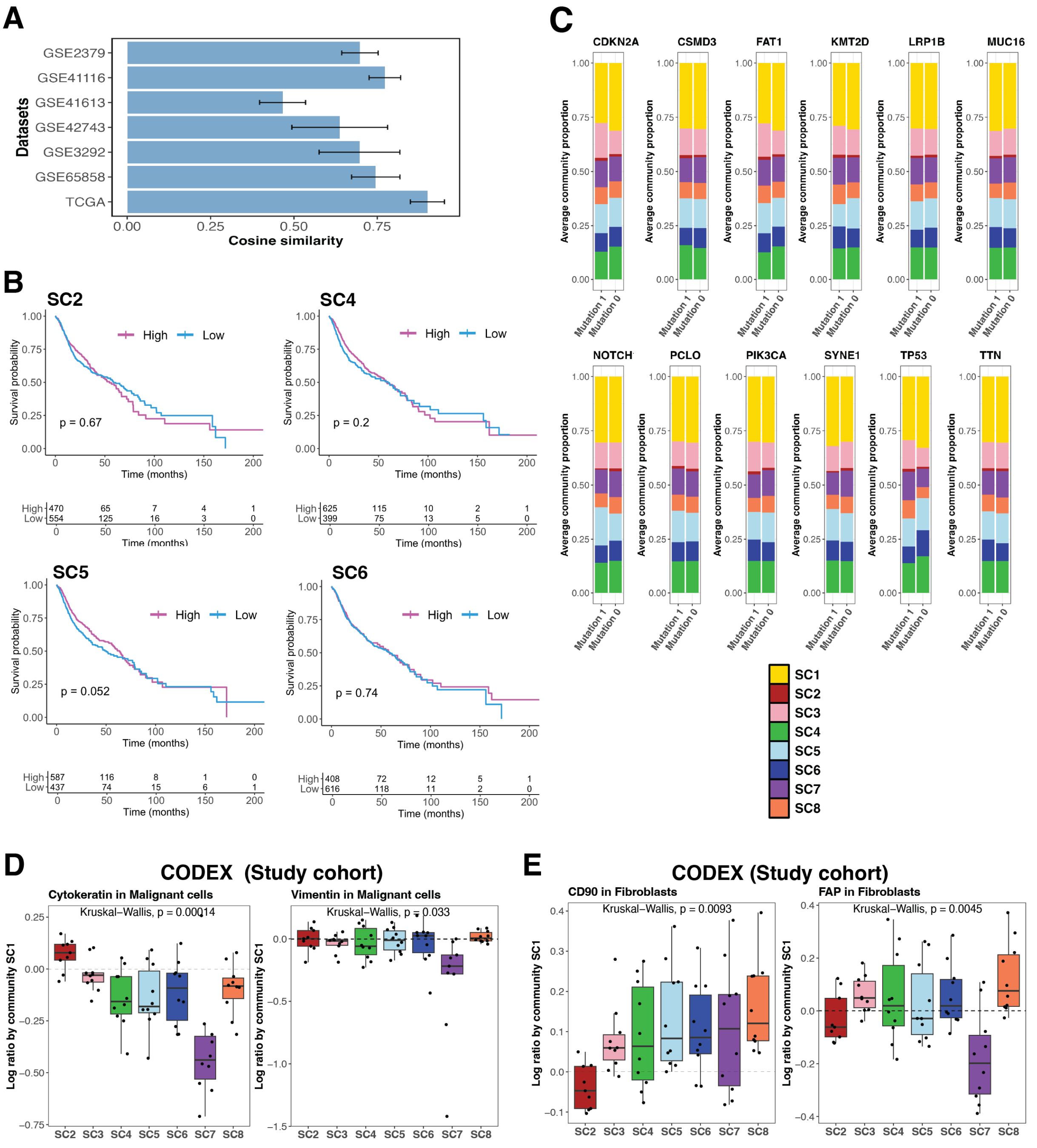


**Figure S7. Additional results of applying MONTAGE to deconvolve the public bulk RNA-seq datasets.**

**(A)** Similarity scores between deconvolved spatial community (SC) compositions from public bulk RNA-seq datasets and SC compositions identified from CODEX data in the study cohort.

**(B)** Kaplan-Meier survival curve of other non-prognostic MONTAGE deconvolved public bulk RNA-seq SCs.

**(C)** Barplot showing SC proportions between mutation status of genes with mutation frequency higher than 15% in TCGA HNSCC mutation data.

**(D)** Ratios of Cytokeratin and Vimentin protein expressions in the malignant cells against malignant cells in SC1 (tumor enriched) across all other SCs in the CODEX data from the study cohort.

**(E)** Ratios of CD90 and FAP protein expressions in the fibroblast cells against fibroblast cells in SC1 (tumor enriched) across all other SCs in the CODEX data from the study cohort.

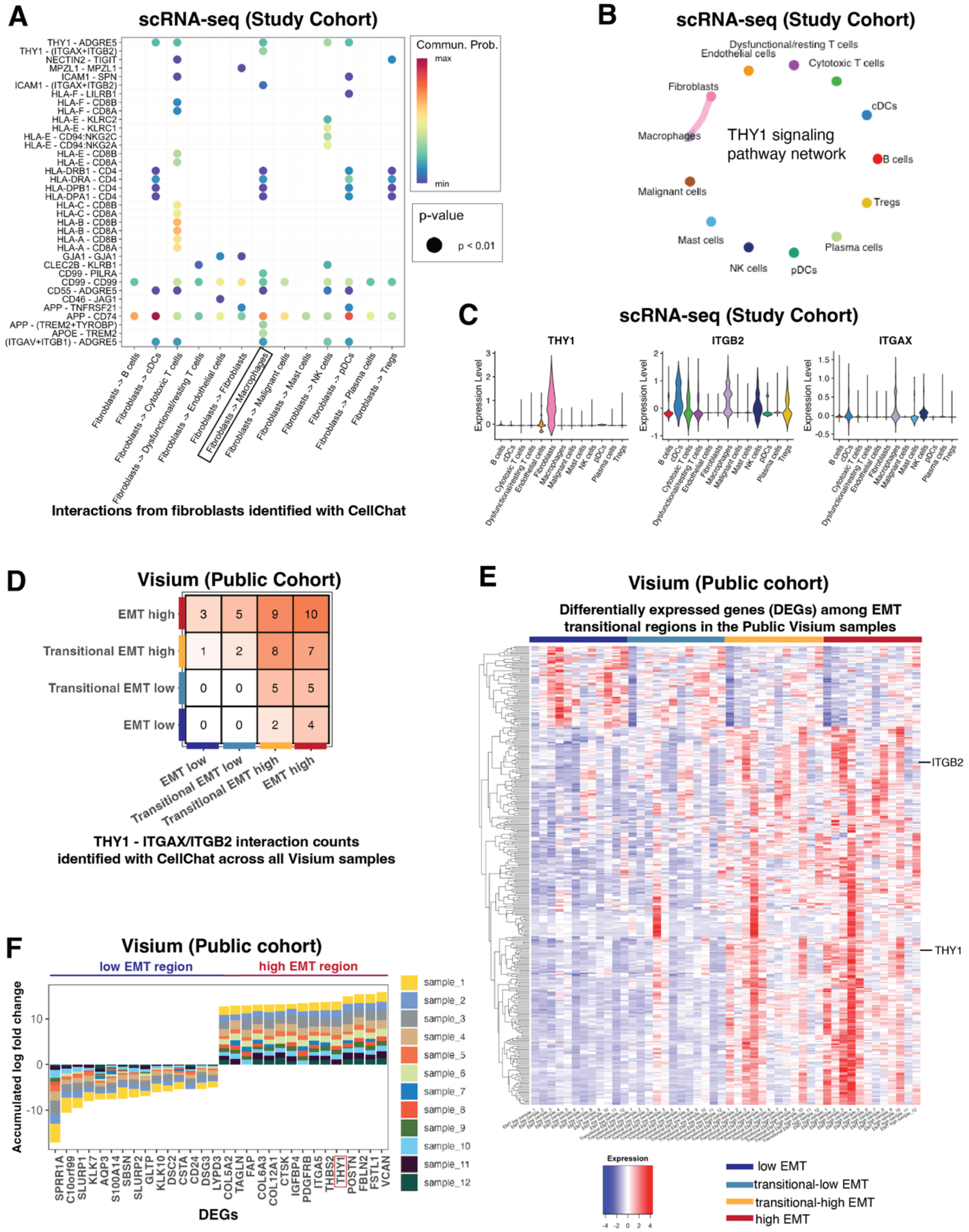
A
scRNA-seq (Study Cohort)
THY1 - ADGRE5
THY1 - (ITGAX+ITGB2)
NECTIN2 - TIGIT
MPZL1 - MPZL1
ICAM1 - SPN
ICAM1 - (ITGAX+ITGB2)
HLA-F - LILRB1
HLA-F - CD8B
HLA-F - CD8A
HLA-E - KLRC2
HLA-E - KLRC1
HLA-E - CD94:NKG2C
HLA-E - CD94:NKG2A
HLA-E - CD8B
HLA-E - CD8A
HLA-DRB1 - CD4
HLA-DRA - CD4
HLA-DPB1 - CD4
HLA-DPA1 - CD4
HLA-C - CD8B
HLA-C - CD8A
HLA-B - CD8B
HLA-B - CD8A
HLA-A - CD8B
HLA-A - CD8A
GJA1 - GJA1
CLEC2B - KLRB1
CD99 - PILRA
CD99 - CD99
CD55 - ADGRE5
CD46 - JAG1
APP - TNFRSF21
APP - CD74
APP - (TREM2+TYROBP)
APOE - TREM2
(ITGAV+ITGB1) - ADGRE5
Fibroblasts -> B cells
Fibroblasts -> cDCs
Fibroblasts -> Cytotoxic T cells
Fibroblasts -> Dysfunctional/resting T cells
Fibroblasts -> Endothelial cells
Fibroblasts -> Fibroblasts
Fibroblasts -> Macrophages
Fibroblasts -> Malignant cells
Fibroblasts -> Mast cells
Fibroblasts -> NK cells
Fibroblasts -> pDCs
Fibroblasts -> Plasma cells
Fibroblasts -> Tregs
Commun. Prob.
max
min
p-value
p < 0.01
Interactions from fibroblasts identified with CellChat
B
scRNA-seq (Study Cohort)
Dysfunctional/resting T cells
Endothelial cells
Cytotoxic T cells
Fibroblasts
cDCs
Macrophages
THY1 signaling
pathway network
B cells
Malignant cells
Tregs
Mast cells
Plasma cells
NK cells
pDCs
C
scRNA-seq (Study Cohort)
THY1
ITGB2
ITGAX
Expression Level
B cells
cDCs
Cytotoxic T cells
Dysfunctional/resting T cells
Endothelial cells
Fibroblasts
Macrophages
Malignant cells
Mast cells
NK cells
pDCs
Plasma cells
Tregs
D
Visium (Public Cohort)
EMT high
Transitional EMT high
Transitional EMT low
EMT low
3 5 9 10
1 2 8 7
0 0 5 5
0 0 2 4
EMT low
Transitional EMT low
Transitional EMT high
EMT high
THY1 - ITGAX/ITGB2 interaction counts
identified with CellChat across all Visium samples
E
Visium (Public cohort)
Differentially expressed genes (DEGs) among EMT
transitional regions in the Public Visium samples
ITGB2
THY1
Expression
-4 -2 0 2 4
low EMT
transitional-low EMT
transitional-high EMT
high EMT
F
Visium (Public cohort)
low EMT region
high EMT region
Accumulated log fold change
10
0
-10
SPRR1A
C10orf99
SLURP1
KLK7
AQP3
S100A14
SBSN
SLURP2
GLTP
KLK10
DSC2
CSTA
CD24
DSG3
LYPD3
COL5A2
TAGLN
FAP
COL6A3
COL12A1
CTSK
IGFBP4
PDGFRB
ITGA5
THBS2
THY1
POSTN
FBLN2
FSTL1
VCAN
DEGs
sample_1
sample_2
sample_3
sample_4
sample_5
sample_6
sample_7
sample_8
sample_9
sample_10
sample_11
sample_12

**Figure S8. Additional results of CD90 (coded by THY1 gene) and integrin crosstalk mediating fibroblast- and macrophage-enrichment.**

**(A)** CellChat identified ligand-receptor interactions from fibroblast cells to other cell types in the scRNA-seq data of the study cohort.

**(B)** CellChat identified that THY1 signaling pathway is only significantly connected between fibroblast cells and macrophages in the scRNA-seq data from the study cohort.

**(C)** THY1, ITGAX, ITGB2 expressions across cell types in the study cohort scRNA-seq data.

**(D)** Numbers of significant THY1-ITGAX/ITGB2 interactions in different EMT regions across the 12 samples in the public Visium dataset identified by CellChat.

**(E)** Heatmap displaying differentially expressed genes (DEGs) among the EMT regions in the public Visium dataset.

**(F)** Top 15 DEGs between the low and high EMT regions in the public Visium dataset.

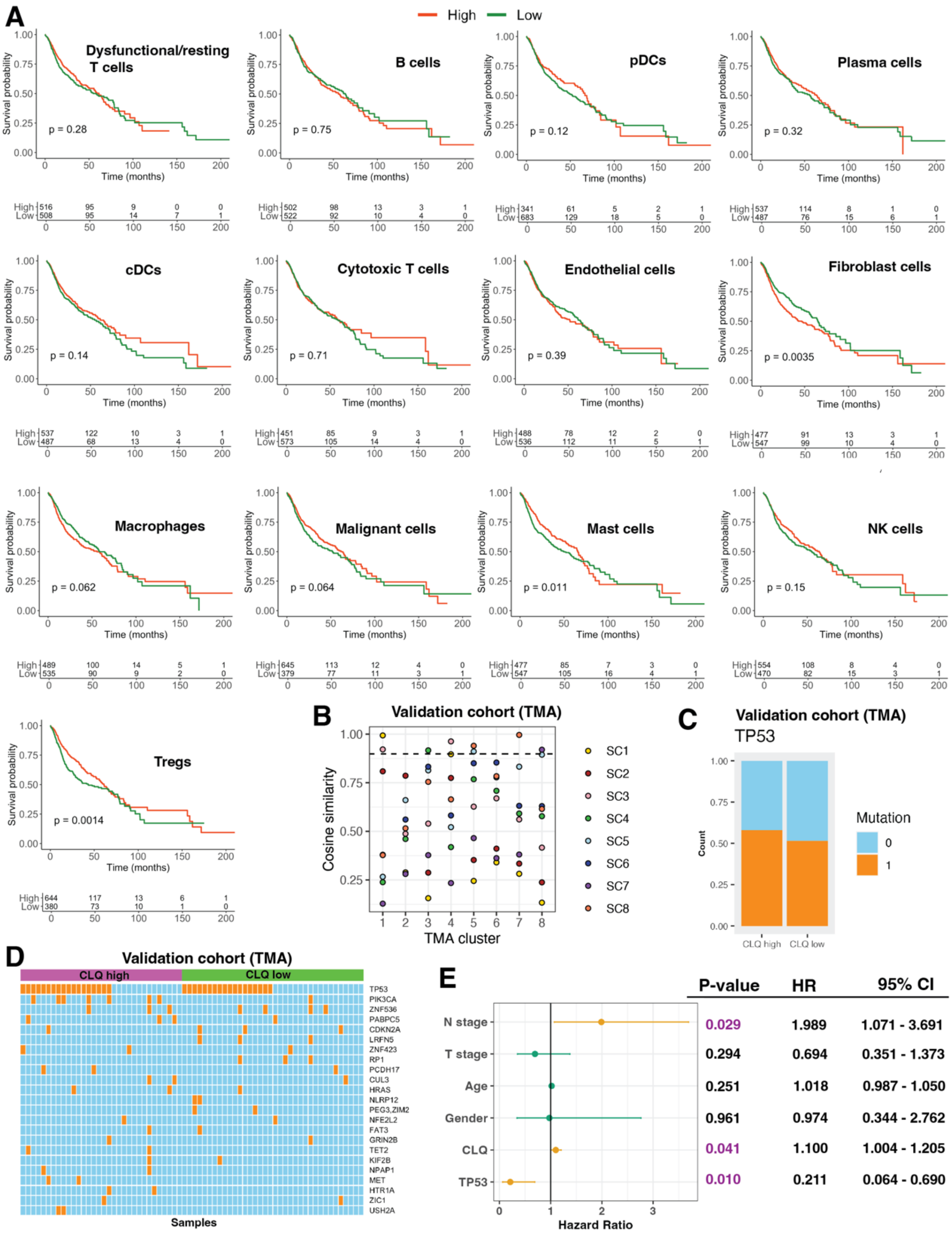
A
High
Low
Dysfunctional/resting T cells
p = 0.28
B cells
p = 0.75
pDCs
p = 0.12
Plasma cells
p = 0.32
cDCs
p = 0.14
Cytotoxic T cells
p = 0.71
Endothelial cells
p = 0.39
Fibroblast cells
p = 0.0035
Macrophages
p = 0.062
Malignant cells
p = 0.064
Mast cells
p = 0.011
NK cells
p = 0.15
Tregs
p = 0.0014
Survival probability
Time (months)
B
Validation cohort (TMA)
Cosine similarity
TMA cluster
SC1
SC2
SC3
SC4
SC5
SC6
SC7
SC8
C
Validation cohort (TMA)
TP53
Count
Mutation
0
1
CLQ high
CLQ low
D
Validation cohort (TMA)
CLQ high
CLQ low
TP53
PIK3CA
ZNF536
PABPC5
CDKN2A
LRFN5
ZNF423
RP1
PCDH17
CUL3
HRAS
NLRP12
PEG3,ZIM2
NFE2L2
FAT3
GRIN2B
TET2
KIF2B
NPAP1
MET
HTR1A
ZIC1
USH2A
Samples
E
P-value
HR
95% CI
N stage 0.029 1.989 1.071 - 3.691
T stage 0.294 0.694 0.351 - 1.373
Age 0.251 1.018 0.987 - 1.050
Gender 0.961 0.974 0.344 - 2.762
CLQ 0.041 1.100 1.004 - 1.205
TP53 0.010 0.211 0.064 - 0.690
Hazard Ratio

**Figure S9. Additional results of public bulk RNA-seq deconvolution and validation cohort TMA analysis.**
**(A)** The Kaplan-Meier curves of deconvolved public bulk RNA-seq cell type composition survival analysis.
**(B)** Comparing the spatial communities (SCs) identified on the CODEX data of the TMA cores with the SCs identified on the whole-slide CODEX data in the study cohort.
**(C)** Mutation status of TP53 between fibroblast and macrophage co-localization (CLQ) high and low groups of the validation TMA cohort.
**(D)** Heatmap showing the mutation status of major mutations (mutation frequency > 3%) between fibroblast and macrophage co-localization (CLQ) high and low groups of the validation TMA cohort.
**(E)** The Cox proportional hazard multi-variate survival analysis of fibroblast and macrophage co-localization (CLQ) and clinical factors of the validation TMA cohort. HR: hazard ratio. CI: Confidence Interval.

### Supplementary Tables

| Sample ID | Patient ID | Tissue type | TMN Stage | P16 | Dataset |
|---|---|---|---|---|---|
| SCC7233 | SCC7233 | Primary tumor | T3 /N3b/Mx | Negative | CODEX |
| SCC7233B | SCC7233 | Metastatic lymph node | T3 /N3b/Mx | Negative | CODEX |
| SCC7238 | SCC7238 | Primary tumor | T4a/N2b/Mx | NA | scRNA-seq&CODEX |
| SCC7239 | SCC7239 | Primary tumor | NA | NA | scRNA-seq |
| SCC7239B | SCC7239 | Metastatic lymph node | NA | NA | scRNA-seq |
| SCC7240 | SCC7240 | Primary tumor | T3/N2b/Mx | Negative | scRNA-seq&CODEX |
| SCC7240B | SCC7240 | Metastatic lymph node | T3/N2b/Mx | Negative | scRNA-seq&CODEX |
| SCC7267 | SCC7267 | Primary tumor | T4a/N3/Mx | Negative | CODEX |
| SCC7267B | SCC7267 | Metastatic lymph node | T4a/N3/Mx | Negative | scRNA-seq&CODEX |
| SCC7268 | SCC7268 | Primary tumor | T3/N0/Mx | NA | scRNA-seq&CODEX |
| SCC7275 | SCC7275 | Primary tumor | T4a/ N0/Mx | Negative | scRNA-seq&CODEX |
| SCC7276B | SCC7276 | Metastatic lymph node | T2/N2/Mx | Positive | scRNA-seq&CODEX |

**Supplementary Table S1: Patient clinical information of the HNSCC study cohort.**

| Dataset | Sample size | Survival outcome | Mutation data | Stage information | HPV data | Age | Gender |
|---|---|---|---|---|---|---|---|
| TCGA | n=520 | Yes | Yes | Yes | Yes | Yes | Yes |
| GSE65858 | n=253 | Yes | No | No | No | No | No |
| GSE41613 | n=97 | Yes | No | No | All HPV negative | No | No |
| GSE42743 | n=74 | Yes | No | No | No | No | No |
| GSE41116 | n=43 | Yes | No | No | Yes | Yes | Yes |
| GSE3292 | n=32 | Yes | No | Yes | Yes | Yes | Yes |
| GSE2379 | n=9 | Yes | No | Yes | No | No | No |

**Supplementary Table S3: Characterization of public HNSCC bulk RNA-seq datasets.**
